# Effects of different slow paced breathing regimes on cerebrospinal fluid (CSF) oscillations

**DOI:** 10.64898/2025.12.24.696312

**Authors:** Padideh Nasseri, Makaila Banks, Albert Y. Li, Laura D. Lewis, Hyun Joo Yoo, Mara Mather

## Abstract

Cerebrospinal fluid (CSF) transport is crucial for clearing waste from the brain and is largely driven by blood vessel oscillatory dynamics. During non-REM sleep, large oscillations occur in the infraslow range around 0.02-0.03 Hz, where noradrenergically driven vasomotion, blood volume and CSF flow are tightly coupled. Whether an infraslow oscillation induced by respiration can engage comparable dynamics during wakefulness is unknown. Utilizing a fast neuroimaging technique, we investigated the effects of four different slow paced respiratory conditions on CSF oscillations in 35 healthy adults (aged 18-30): paced breathing at 0.09-0.11 Hz and 0.05-0.07 Hz, and paced sighing at 0.05-0.07 Hz and 0.02-0.03 Hz. All four conditions increased respiratory volume, low-frequency respiratory power, CSF oscillatory power and BOLD global signal amplitude relative to rest, and CSF power did not differ among them. However, the infraslow slow sighing produced significantly higher CSF-BOLD coupling than all the other conditions, together with the largest global signal amplitude, an index associated with reduced arousal. This effect of infraslow sighing was not linked with respiratory power, as paced breathing at 0.05-0.07 Hz had the greatest respiratory power yet the weakest CSF-BOLD coupling, falling below rest. Cardiac spectral power was also lower during infraslow sighing than during any other breathing manipulation. Stimulating oscillatory dynamics via respiration during wakefulness therefore may recruit neurovascular-CSF coupling due to the same infraslow-specific dynamics that are believed to drive sleep-dependent waste clearance. Our study provides evidence that slow paced breathing increases CSF oscillations during wakefulness. Furthermore, we found that sighing at the frequency of the infraslow band linked to glymphatic clearance increases the coupling between blood and CSF dynamics.

**Highlights:**

- Slow paced respiration and sighing both increased CSF oscillatory power
- Infraslow paced sighing led the strongest dynamics between CSF and BOLD
- Breathing and sighing manipulations lowered the BOLD signal proxy of arousal

## Introduction

Efficient waste removal plays a vital role in maintaining organ homeostasis. In the central nervous system, cerebrospinal fluid (CSF) serves as a key medium for solute transport and the clearance of metabolic waste from brain tissue, including beta amyloid and tau (Benveniste et al., 2018; Iliff et al., 2013; Xie et al., 2013). Impaired clearance is observed in healthy aging and, more markedly, in age-related neurodegenerative disorders including Alzheimer’s disease (Li et al., 2022; Simon & Iliff, 2016; Zhou et al., 2020), making the physiological drivers of CSF movement important to understand.

CSF movement is constrained by the Monro-Kellie principle that the combined volume of brain tissue, blood and CSF are constant (Mokri, 2001). Because the cranium is rigid and brain tissue is not easily compressed, any increase in intracranial blood volume must be offset by a compensatory displacement of CSF, and any decrease permits a net influx. CSF flow therefore depends mechanically on oscillations in cerebral blood volume. Human imaging has confirmed this relationship, showing an inverse coupling between cerebral blood volume and CSF flow (Fultz et al., 2019; Picchioni et al., 2022; Väyrynen et al., 2026; Yang et al., 2022; Zimmermann et al., 2025). Importantly, because it is the change in cerebral blood volume that affects CSF flow, it is the rate of change in the BOLD signal, or the negative BOLD derivative, that is associated with CSF inflow, rather than the instantaneous amplitude of BOLD signal (Fultz et al., 2019; Zimmermann et al., 2025).

Insofar as CSF movement is driven by oscillatory changes in vascular volume, the frequency and amplitude of those changes should determine how much CSF flows in or out of the cranium. Different aspects of physiology cause oscillations in vascular volume at different frequencies. Cardiac pulsations are the fastest and were implicated in CSF oscillations early on. For instance, in rodents, increasing arterial pulsatility by administering dobutamine, an adrenergic agonist, increased CSF penetration into the brain’s parenchyma whereas reducing arterial pulsatility through internal carotid artery ligation diminished CSF penetration and CSF–interstitial fluid exchange (Iliff et al., 2013). Human MRI likewise shows cardiac-locked CSF pulsations (Feinberg & Mark, 1987; Mascalchi et al., 1988). However, cardiac pulsations only change brain vessel diameter by less than 1% in rodents (Iliff et al., 2013). In contrast, the periodic constriction and dilation of cerebral arteries below 0.1 Hz, known as slow vasomotion, produces much larger diameter changes of around 10-15%, corresponding to a ≈20-30% increase in vascular volume (Drew et al., 2011; Nedergaard, 2026; Veluw et al., 2020a). Indeed, in awake mice, both spontaneous vasomotion at approximately 0.1 Hz and evoked vasomotion at 0.05 Hz have been shown to drive CSF flow, with greater vasomotion amplitudes associated with higher clearance rates (Veluw et al., 2020b). Visual stimulation at 0.03 Hz and 0.1 Hz in humans induces CSF movement in the fourth ventricle and visual cortex, indicating that neurovascular dynamics evoked at a chosen frequency can be used to move CSF (Hirschler et al., 2025; Williams et al., 2023).

Sleep research suggests that infraslow vascular oscillations (peaking at around 0.02-0.03 Hz) may be the most consequential for CSF movement. During sleep, levels of noradrenaline oscillate in the range of 0.02 Hz in rodents (Kjaerby et al., 2026; Nedergaard, 2026; Osorio-Forero et al., 2021), with similar oscillatory dynamics seen in levels of acetylcholine, serotonin and dopamine (Kjaerby et al., 2026). In rodents, use of optogenetic activation and pharmacological manipulation sdemonstrated that locus-coeruleus-mediated noradrenaline release drives vasomotion during non-rapid eye movement (NREM) sleep, which in turn drives CSF inflow into the brain (Haugland et al., 2025). In humans during sleep, the infraslow power of BOLD fMRI increases significantly, with a peak at 0.03 Hz, indicating an increase in infraslow vasomotion at around the same frequency range as in rodents (Horovitz et al., 2008; Väyrynen et al., 2026). Recently, this infraslow band has been proposed as the central mechanical driver of glymphatic clearance (Hauglund & Nedergaard, 2026; Nedergaard, 2026).

Two features of the literature motivated our study. First, the infraslow oscillations in CSF are not unique to sleep. Coupling between infraslow CSF flow and cerebrovascular oscillations is detectable during wakefulness (H.-C. (Shawn) Yang et al., 2022) and sleep-deprived but awake participants show waves of increased cortical CSF flow that are synchronized with attentional lapses, pupil dilation and infraslow neurovascular dynamics (Z. Yang et al., 2025). Thus, it seems feasible to engage infraslow dynamics during wakefulness. Second, this infraslow band is driven by noradrenergic oscillations, and lower tonic noradrenergic levels facilitate CSF movement potentially because high noradrenergic tone limits oscillatory dynamics (Xie et al., 2013; Haugland et al., 2025). A manipulation that both stimulates an infraslow vascular oscillation and lowers arousal could potentially act on CSF dynamics through two convergent routes.

Respiration is a strong candidate for such a manipulation because it is the only major driver of cerebral blood volume that is under voluntary control. Interestingly, respiration is entrained to the evolutionarily conserved sleep-enhanced infraslow rhythms that have been identified across seven species, including humans (Bergel et al., 2026). Respiratory-frequency pulsations are also clearly visible in the CSF signal during wakefulness (Fultz et al., 2019). Forced inspiration induces an upward movement of CSF toward the head, whereas forced expiration produces a downward movement of CSF (Aktas et al., 2019; Dreha-Kulaczewski et al., 2017), with abdominal breathing being associated with higher CSF movement compared to thoracic breathing (Aktas et al., 2019; Yildiz et al., 2017). Furthermore, forced breathing shifted the dominant frequency of both CSF and venous flow spectra towards the respiratory component and increased the magnitude of CSF flow (Kollmeier et al., 2022). Meditation practices involving slow, deep breathing, such as yogic breathing and Seokmun Hoheup meditation, have similarly been shown to increase CSF flow (Lim et al., 2025; Yildiz et al., 2022). Slow paced breathing also amplifies cardiovascular oscillations through baroreceptor resonance near 0.1 Hz (Lehrer & Gevirtz, 2014; Nuckowska et al., 2019), and we have previously shown that participants performing slow paced breathing exhibit higher BOLD spectral power than controls, with the peak power at the paced breathing frequency (Nashiro et al., 2021). Respiration further engages the locus coeruleus-norepinephrine system directly (Yackle et al., 2017), and slow breathing reduces peripheral noradrenergic activity (Oneda et al., 2010), providing the arousal/noradrenaline reducing route described above.

A challenge, however, is that to entrain the infraslow band of 0.02-0.03 Hz using breathing would require breath cycles of 30-50 s, which is not sustainable. Paced sighing provides an effective alternative. Paced sighing at lower frequencies has been shown to create strong oscillations in the cardiovascular system (B. Vaschillo & Vaschillo, 2020; E. G. Vaschillo et al., 2015). Sighing, here described as deep inhales followed by longer exhales, is linked to resetting of respiratory rate, transition between autonomic states such as the transition from wakefulness to NREM sleep, and psychological relief (Ramirez, 2014; Severs et al., 2022; Vlemincx et al., 2016). Similar to paced breathing, paced sighing entrains vascular tone and arterial elasticity (B. Vaschillo & Vaschillo, 2020). Sighing therefore allows a way to produce a vascular oscillation at the infraslow band that rodent and human sleep work has identified as the dominant driver of glymphatic transport.

Here, we use fast neuroimaging to compare CSF oscillations across four paced respiratory conditions spanning three frequency ranges: 0.09-0.11 Hz, 0.05-0.07 Hz, and 0.02-0.03 Hz, with the last achieved through paced sighing rather than paced breathing, and the middle band including a comparison of paced breathing and paced sighing at the same frequency. Critically, given that the infraslow band is where noradrenergically driven vasomotion, cerebral blood volume and CSF flow are most tightly coupled during sleep (Hauglund et al., 2025; Helakari et al., 2022; Väyrynen et al., 2026), we hypothesized that respiratory oscillations at the 0.02-0.03 Hz frequency would lead to the strongest coupling between CSF oscillations and the negative derivative of the BOLD signal.

## Method

### Participants

We calculated the sample size for this project using G*Power (Version 3.1.9; Faul et al., 2007). Prior recent human studies investigating respiration effects on CSF dynamics generally included relatively small within-subject samples, typically ranging from 8-26 participants (Dreha-Kulaczewski et al., 2017; Kollmeier et al., 2022; Nair et al., 2024; Yildiz et al., 2022). We set the power of our study to detect medium to large effects. Our sample size estimation indicated that 31 participants would provide 95% power to detect a within-subject effect size of f=0.25 in a repeated measures analysis of variance (ANOVA), with an alpha level of 0.05. To accommodate potential exclusions and dropouts, we targeted a sample size of 39 participants.

Eligible participants were healthy adults aged 18–35 years who were proficient in English and had normal or corrected-to-normal vision (contact lenses only) and normal hearing. Individuals were excluded if they had a major medical, neurological, or psychiatric illness; a condition that would interfere with paced breathing or preclude MRI scanning (e.g., coronary artery disease, angina, or a cardiac pacemaker); were currently practicing relaxation, biofeedback, or breathing training; or were taking psychoactive medications other than antidepressants or antianxiety medications. Participants taking antidepressants or antianxiety medications were eligible only if their treatment had remained stable for at least 3 months. Prospective participants were screened for eligibility, and those who met all inclusion criteria provided written informed consent under a protocol approved by the University of Southern California (USC) Institutional Review Board.

Participants attended two sessions and received $20 per hour for each visit. Each visit lasted approximately 1.5 hours. Thirty-nine participants were initially recruited. However, three dropped out after the first session, and one participant couldn’t complete the MRI session due to newly recognized claustrophobia. Therefore, 35 participants (18 female), aged 18 to 30 years (M_age_= 22.03, SD_age_=3.01), completed the study and were included in the cross-correlation and CSF spectral frequency analyses. One participant’s physiological data was not recorded due to technical problems, and another participant’s physiological data during the resting state scan was interrupted due to a technical error. Therefore, both were excluded from the respiration and cardiac data analysis. See Fig. 1 for the study flow chart, and Table 1 for the study group characteristics (N =35).

**Fig. 1.**
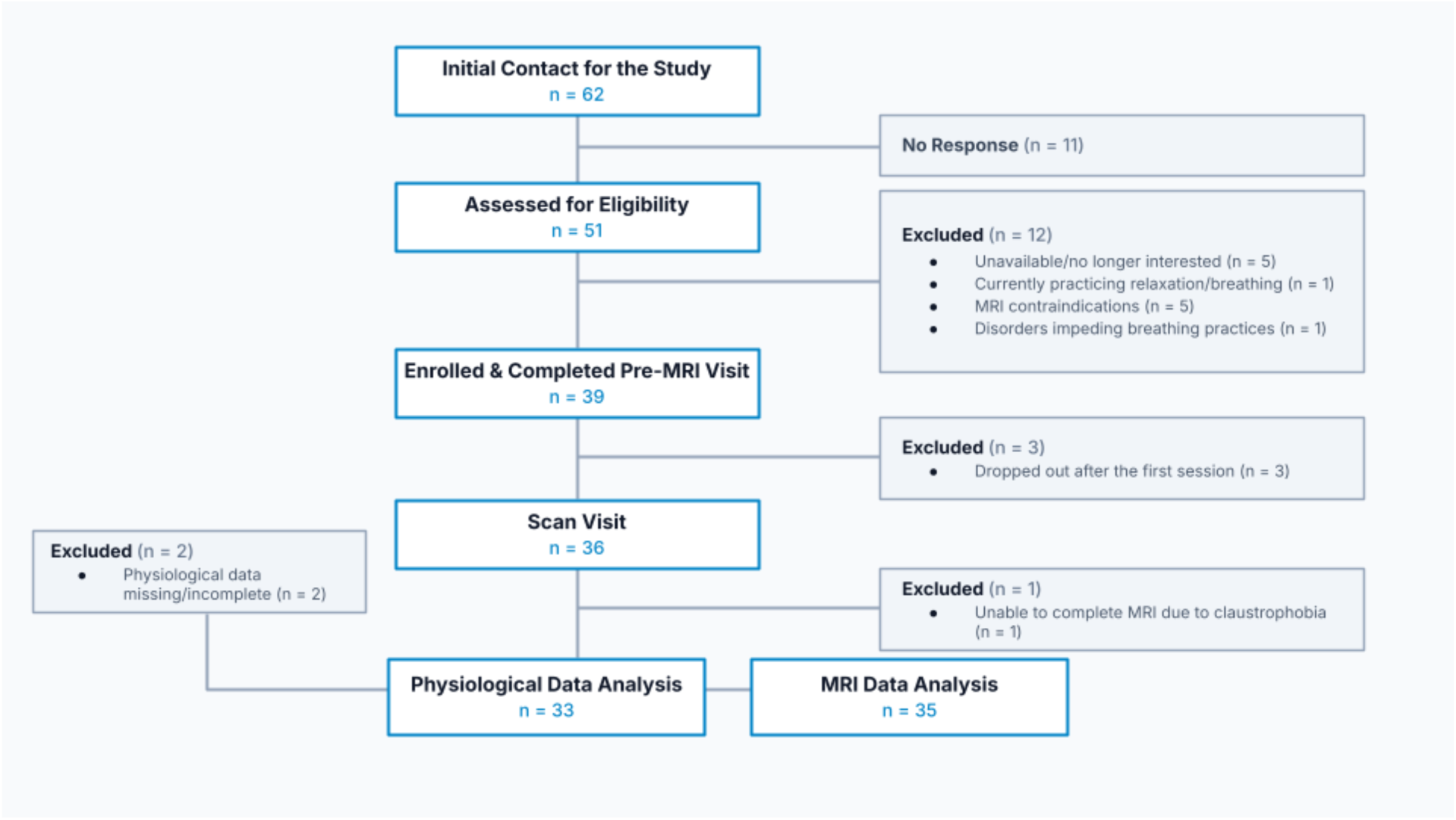
Study flow chart showing the number of participants enrolled and excluded at each stage of study implementation and data analysis. For additional details on the exclusion criteria, see the main text.

**Table 1.** Study group characteristics.

| N | Age range | Age in years<br>Mean (SD) | Sex | Race | Ethnicity |
| --- | --- | --- | --- | --- | --- |
| 35 | 18-30 | 22.03 (3.01) | F, 18 | 13 Asian | 6 Hispanic or Latino |
|  |  |  |  | 16 White |  |
|  |  |  | M, 17 | 1 More than one race | 29 Not |
|  |  |  |  | 5 Other/Prefer not to state | Hispanic or Latino |

### Breathing conditions

Four different paced breathing exercises were used. These included paced breathing at a frequency in the 0.09-0.11 Hz range (1 breath cycle per 9-11 seconds with similar inhalation and exhalation duration) and in the 0.05-0.07 Hz range (1 breath cycle per 14-20 seconds with similar inhalation and exhalation duration) and paced sighing in the 0.05-0.07 Hz range (one sigh every 14-20 seconds) and in the 0.02-0.03 Hz range (one sigh every 33-50 seconds). During the paced breathing conditions, inhalation and exhalation were equal in duration (e.g., 5 seconds of inhalation followed by 5 seconds of exhalation). In our study, paced sighing conditions were defined as alternating periods of normal breathing and sighing. A sigh was characterized by a brief inhalation (1 second) followed by a prolonged exhalation (3 seconds).

### Study procedure

Participants attended two sessions. In the first session, they were introduced to paced breathing and paced sighing. To account for interindividual variability in endogenous cardiovascular dynamics, we individualized the breathing frequencies based on dominant peaks observed in each participant’s resting cardiac spectrum within frequency ranges of interest. Based on principles of resonance (Lehrer et al., 2000; Schwerdtfeger et al., 2020; E. Vaschillo et al., 2002), pacing breathing at an intrinsic cardiac oscillatory frequency should amplify the cardiac oscillations at that frequency, and thereby, via the Monro-Kellie principle, also amplify CSF oscillations.

In order to identify each participant’s individual cardiac oscillatory peaks within targeted frequency ranges, they completed 5 minutes of paced breathing at 0.25 Hz (4 seconds per breath cycle). This specific rate, based on a prior study (Sakakibara et al., 2020), was chosen to control the frequency of breathing, allowing the peaks around the participant’s baroreflex frequency and in the very low frequency (VLF) range to be more detectable. For this assessment of personalized paces, participants wore an ear sensor to measure their pulse, which was connected to a laptop running emWave Pro software. Participants were instructed to breathe in through the nose and out through the mouth in sync with a visual cue on the laptop display. We analyzed the pulse data using Kubios HRV Premium 3.1 software (Tarvainen et al., 2014) and calculated the highest amplitude peaks within 0.09-0.11 Hz, 0.05-0.07 Hz, and 0.02-0.03 Hz ranges on the spectral graph. We used these peak values as the personalized paces for each participant in the study.

After determining each participant’s personalized paces for our ranges of interest, we asked our participants to practice paced breathing and paced sighing at determined personalized paces three times at home before the second lab visit. In the second lab visit, participants underwent MRI scanning sessions, during which they completed 4 breathing manipulation scans (one paced breathing in 0.09-0.11 Hz range, one paced breathing in 0.05-0.07 Hz range, one paced sighing in 0.05-0.07 Hz range, and one paced sighing in 0.02-0.03 Hz range) and one resting state scan.

### MRI data acquisition

MRI data were collected at the University of Southern California David and Dana Dornsife Neuroimaging Center using a Siemens Magnetom Trio 3T MRI scanner equipped with a 32-channel head coil. T1-weighted 3D structural MRI brain scans were acquired using an isotropic magnetization-prepared rapid acquisition gradient echo (MPRAGE) sequence with a TR = 2,300 ms, TE = 2.26 ms, slice thickness = 1.0 mm, flip angle = 9°, field of view = 256 mm, and voxel size = 1.0 × 1.0 × 1.0 mm, collecting 175 volumes over 4 minutes and 44 seconds. Functional MRI scans were performed using a fast fMRI protocol sensitized to detect CSF inflow that consisted of a single-shot gradient echo EPI with multiBand factor = 8, TR = 367 ms, TE = 31.8 ms, slice thickness = 2.5 mm, flip angle = 35°, voxel size = 2.5 × 2.5 × 2.5 mm, FOV = 205mm, number of slices = 40, and no in-plane acceleration.

Each MRI session started with the structural scan for anatomical reference. The functional scans followed the structural scan. For functional scans, the boundary edge of the imaging volume was placed at the fourth ventricle, as identified using structural scans. While fMRI is typically used to detect local oxygenation changes, fast acquisition paradigms can also capture fluid inflow. Fresh fluid entering the edge of the imaging volume exhibits high signal intensity because it has not yet experienced radiofrequency pulses (Ashenagar et al., 2025; Fultz et al., 2019). Positioning the boundary edge of the imaging volume at the fourth ventricle allowed us to capture the CSF signal as it enters the brain, as shown in Fig. 2 and in prior research (Fultz et al., 2019). During resting-state scans, participants were instructed to relax, breathe normally, keep their eyes open, and fixate on a white cross displayed on a gray screen. During the breathing manipulation scans, participants continued to fixate on the cross while following auditory breathing cues. Depending on the condition, the cues instructed participants when to inhale, exhale, or breathe normally. The timing of the cues was personalized based on each participant’s target breathing frequency for that condition (Fig. 3A). For example, in the paced sighing condition at 0.02–0.03 Hz, a participant with a target frequency of 0.025 Hz (one sigh every 40 seconds) breathed normally for most of the 40-second interval. When it was time to sigh, an auditory cue instructed the participant to inhale deeply, followed by another cue to exhale. A third cue then instructed the participant to resume normal breathing until the next scheduled sigh.

**Fig. 2.**
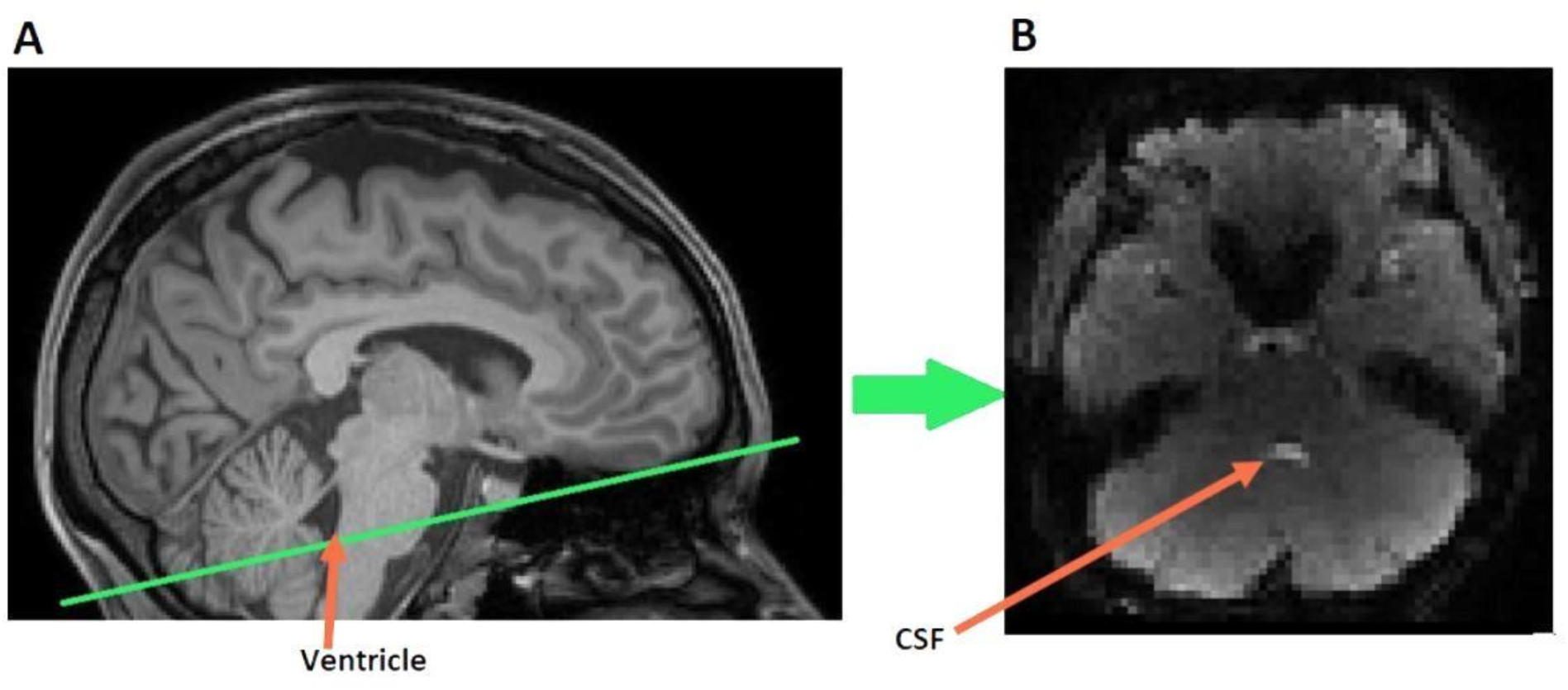
(A) Schematic positioning of fMRI scans. The green line shows positioning of the bottom of the functional image relative to the structural scan. Placing the edge of imaging volume at the fourth ventricle (red arrow) allows capture of CSF inflow. (B) An example image of the bottom slice of functional images. Bright voxels at the fourth ventricle (red arrow) were selected for CSF masks.

**Fig. 3.**
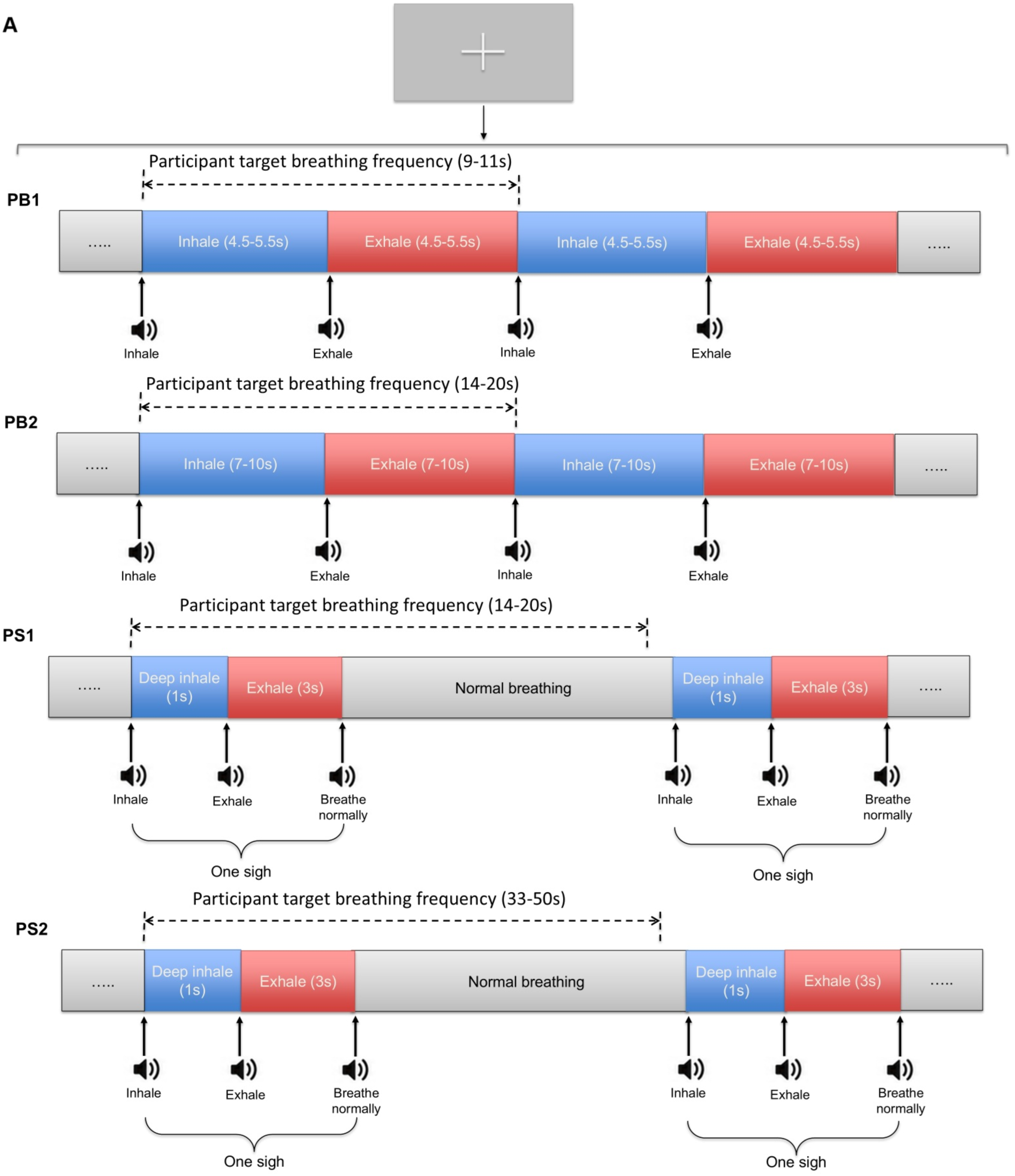
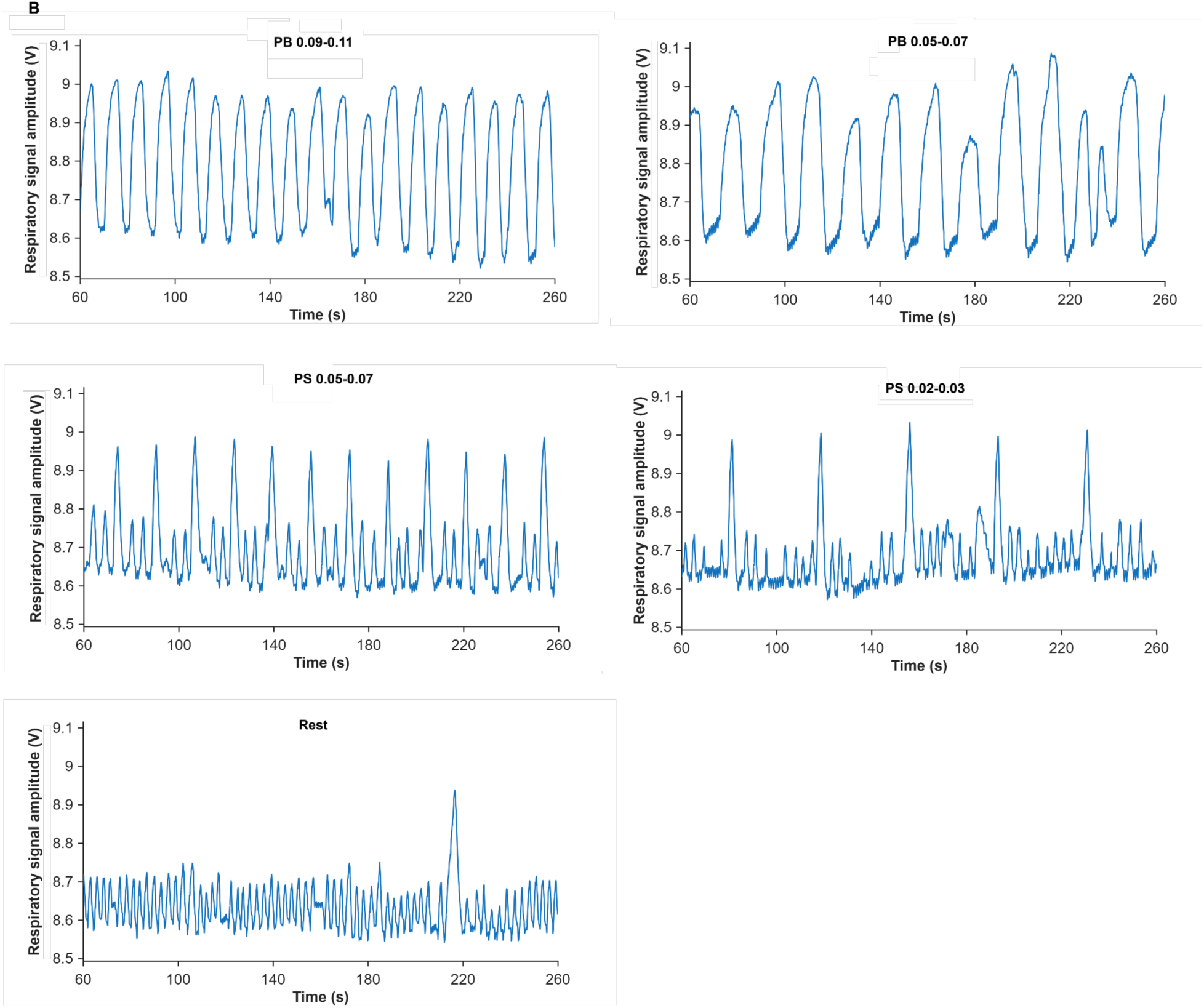
(A) Schematic of the experimental paradigm for paced breathing and paced sighing conditions. Throughout the scans, participants viewed a white fixation cross centered on a gray screen. Auditory cues instructed participants to inhale, exhale, or breathe normally, depending on the condition. Cue timing was individualized based on each participant’s target breathing frequency for each condition. (B) An example of a participant’s respiration signal under different conditions. Note: PB = paced breathing, PS = paced sighing

All participants completed all 4 breathing manipulation scans (paced breathing in 0.09-0.11Hz range, paced breathing in 0.05-0.07 Hz range, paced sighing in 0.05-0.07 Hz range, and paced sighing in 0.02-0.03 Hz range) and resting state scan. Throughout breathing manipulation scans, respiration signals were monitored online using AcqKnowledge® software in real time to facilitate immediate compliance checking. Resting state scans lasted approximately 7 minutes (1144 volumes), while paced breathing and paced sighing scans lasted about 8 minutes (1308 volumes). The order of breathing manipulation scans was randomized across participants. Participants were given short breaks between scans.

### Physiological recording

Throughout functional MRI scans, both photoplethysmogram (PPG) and breathing data were collected using the MR-compatible Biopac MP150 Data Acquisition System. The Biopac Respiratory Effort Transducer (TSD201) was utilized to measure breathing. The electric voltage signal generated by changes in chest circumference was amplified with a gain of 10 times and sampled at 10 kHz using RSP100Cat. The PPG data were collected by using a Nonin Medical 8600FO Pulse Oximeter at a 10 kHz sampling rate.

### Physiological data preprocessing and analysis

Preprocessing and analysis of breathing data were done in MATLAB. The breathing data were downsampled to 1 kHz and subsequently smoothed. We computed respiratory volume (RV) across different conditions using a method developed by Chang et al (2009). This approach involves calculating the sliding-window standard deviation of the respiratory belt waveform (Chang et al., 2009). The RV measure primarily reflects variations in inspiration depth over time and has been shown to correlate with resting-state BOLD signal fluctuations (Chang et al., 2009).

To implement this method, 6-second sliding windows were defined, and the standard deviation of the respiratory signal was calculated within each window. The average of these standard deviations was then computed for each scan and scaled by multiplying the result by 100. The Friedman test was used to compare the RV across different conditions.

Furthermore, to assess the effect of our respiratory modulations on low-frequency (LF) respiratory oscillations, the preprocessed data were band-pass filtered between 0-0.11 Hz and then subjected to Fast Fourier Transform (FFT) to convert the data into the frequency domain. Then, the power spectral density (PSD) was computed. The Friedman test was used to compare the respiratory total and maximum power within the LF range across conditions. The PPG data were downsampled at 1kHz using MATLAB and then were analyzed using Kubios HRV Premium Version 3.1 to compute heart rate, the standard heart rate variability measures of very low frequency heart rate variability (VLF-HRV, 0.00-0.04 Hz), low frequency HRV (LF-HRV, 0.04-0.15 Hz), and high frequency HRV (HF-HRV, 0.15-0.4 Hz).

### Preprocessing of MRI data

The first volume of resting-state scans and the first 165 volumes of breathing manipulation scans were discarded, and non-brain tissue was removed using FSL (https://fsl.fmrib.ox.ac.uk/fsl) BET function. The first 165 volumes of each breathing manipulation scan were discarded to ensure that CSF flow dynamics were measured only after participants had completed at least one full cycle of the breathing manipulation.

For CSF signal extraction, the ArtRepair toolbox (http://cibsr.stanford.edu/tools/human-brain-project/artrepair-software.html) was employed to detect and correct outlier volumes displaying excessive movement within each scan. A threshold of 0.3 mm/TR variation was set to identify outlier volumes. No more than one-third of imaging volumes for each scan were corrected using this method (Table 2). These outlier volumes were repaired using the despike option, which uses a linear interpolation of the immediately preceding and following volumes, while leaving other volumes unaffected. FSL FLIRT was used to align brain-extracted structural images with functional images through a 12-degree of freedom linear affine transformation. The registered structural scans were then overlaid onto the functional images to identify the approximate position of the ventricle/aqueduct. Subsequently, the brightest voxels on the bottom slice of the functional image in the fourth ventricle region were selected to create the ventricle CSF mask (Fig. 1).

**Table 2.**
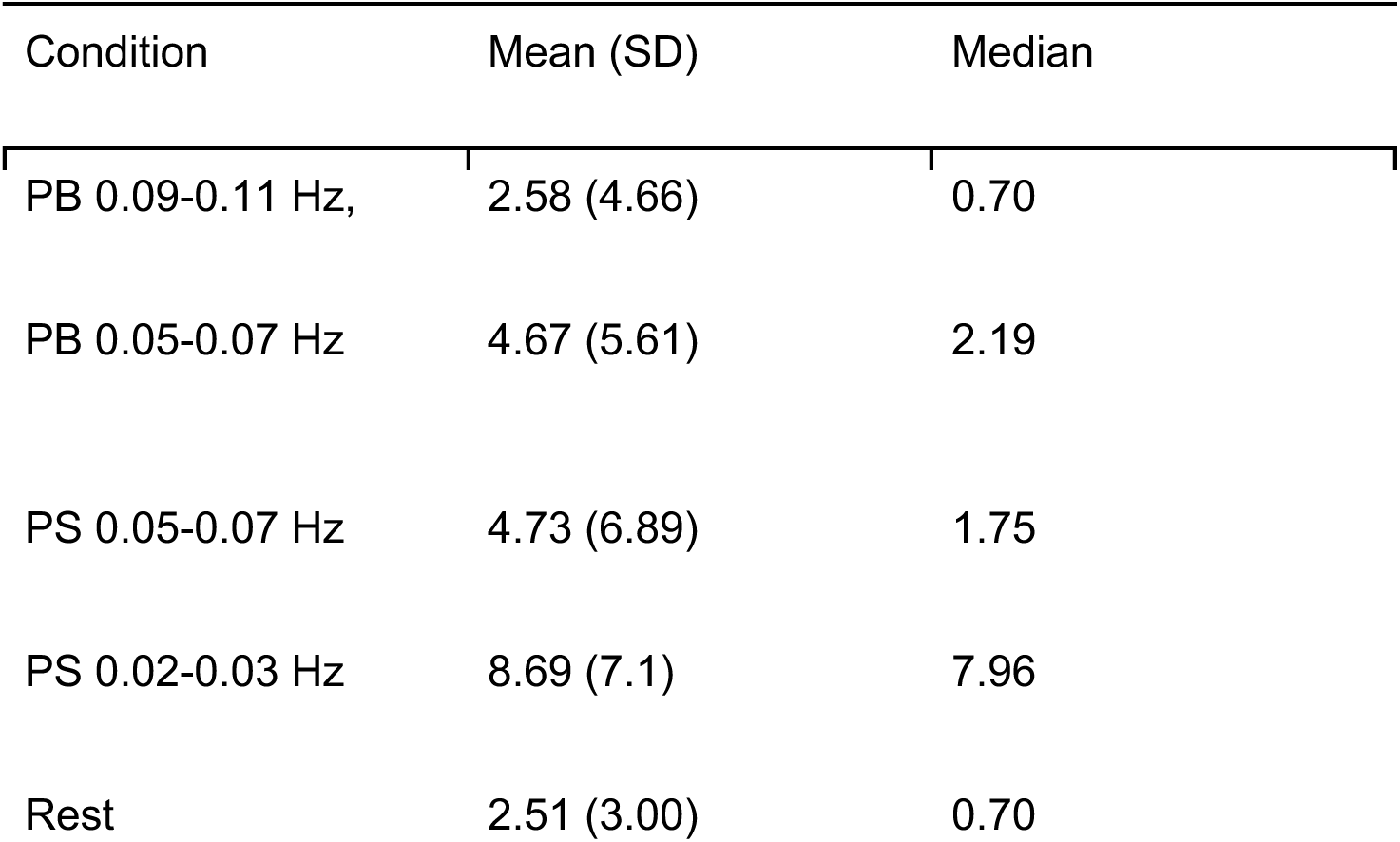
Percentage of corrected volumes by ARTrepair for each scan. PB = paced breathing, PS = paced sighing.

| Condition | Mean (SD) | Median |
| --- | --- | --- |
| PB 0.09-0.11 Hz, | 2.58 (4.66) | 0.70 |
| PB 0.05-0.07 Hz | 4.67 (5.61) | 2.19 |
| PS 0.05-0.07 Hz | 4.73 (6.89) | 1.75 |
| PS 0.02-0.03 Hz | 8.69 (7.1) | 7.96 |
| Rest | 2.51 (3.00) | 0.70 |

For BOLD signal extraction and gray matter BOLD global signal amplitude analysis, raw fMRI images underwent motion correction using FSL MCFLIRT (Jenkinson et al., 2002). FSL FAST was applied to structural images for brain tissue segmentation to generate the gray matter mask. The resulting gray matter mask was registered to functional scans and thresholded at 75%.

### Cross-correlation analysis

CSF time series was extracted from the ventricle ROI of ART-repaired functional scans. BOLD time series were extracted from the gray matter ROI of motion-corrected functional scans. Subsequently, a low-pass temporal filter 0-0.11Hz was applied to both extracted BOLD and CSF time series using MATLAB. Afterward, the time series were detrended. Following detrending, derivatives of the BOLD signal were calculated and then multiplied by -1. Any resulting negative values were subsequently set to zero. Cross-correlation analyses were conducted over lags of -40 to 40 seconds for each subject for each condition.

To establish a threshold for statistical testing and estimate p-values, cluster permutation tests employing the Monte Carlo method with 10,000 permuted randomizations were carried out to build a null distribution for the BOLD–CSF correlation. The p-value for the BOLD–CSF correlations at different lags was obtained by comparing them against the null distribution for each condition. The maximum correlation coefficient from these significant cross-correlation lags was then identified and used to compare the results across conditions using a within-subject repeated measures ANOVA test.

### Spectral frequency analysis of CSF signal

ART-repair corrected functional scans underwent further preprocessing using FSL and Analysis of Functional NeuroImaging (AFNI) (Cox, 1996) tools. These additional steps included temporal despiking, linear detrending, spatial smoothing (full width at half maximum [FWHM] = 3 mm), and global intensity normalization. Then the power spectral density of the data was calculated using FSL’s fslpspec tool. The power spectrum of the fourth ventricle was calculated by using CSF masks over the resulting power spectra. The maximum power among frequencies below 0.11 Hz was calculated for each scan and subject. Then, a within-subject repeated-measures ANOVA was performed to compare the maximum power values below 0.11 Hz among different conditions, as extracted from the CSF power spectra.

### Gray matter BOLD global signal amplitude

The standard deviation of the global signal, known as the global signal amplitude, has been linked to arousal states, with higher values indicating a drowsy state (Wong et al., 2013). To compute this measure, we derived the percent change time series for each voxel by first subtracting the mean value from the preprocessed and motion-corrected scans and then dividing this difference by the mean value. The global mean signal was then determined as the average of these percent change time series across all voxels within the whole-brain gray matter. The standard deviation of this global signal was calculated and referred to as the global signal amplitude. The Friedman test was used to compare the global signal amplitude across different conditions.

## Results

### Breathing modulations change respiration and cardiac metrics

Looking at the RV across conditions showed a significant main effect of condition *χ*^2^(4) = 57.624, *p*<0.001. Post-hoc pairwise comparisons using the Wilcoxon signed-rank tests with FDR correction showed that all breathing manipulation conditions had significantly higher RV than rest (all FDR-corrected p-values < 0.0001; Fig. 4A and Supplemental Table 1).

**Fig. 4.**
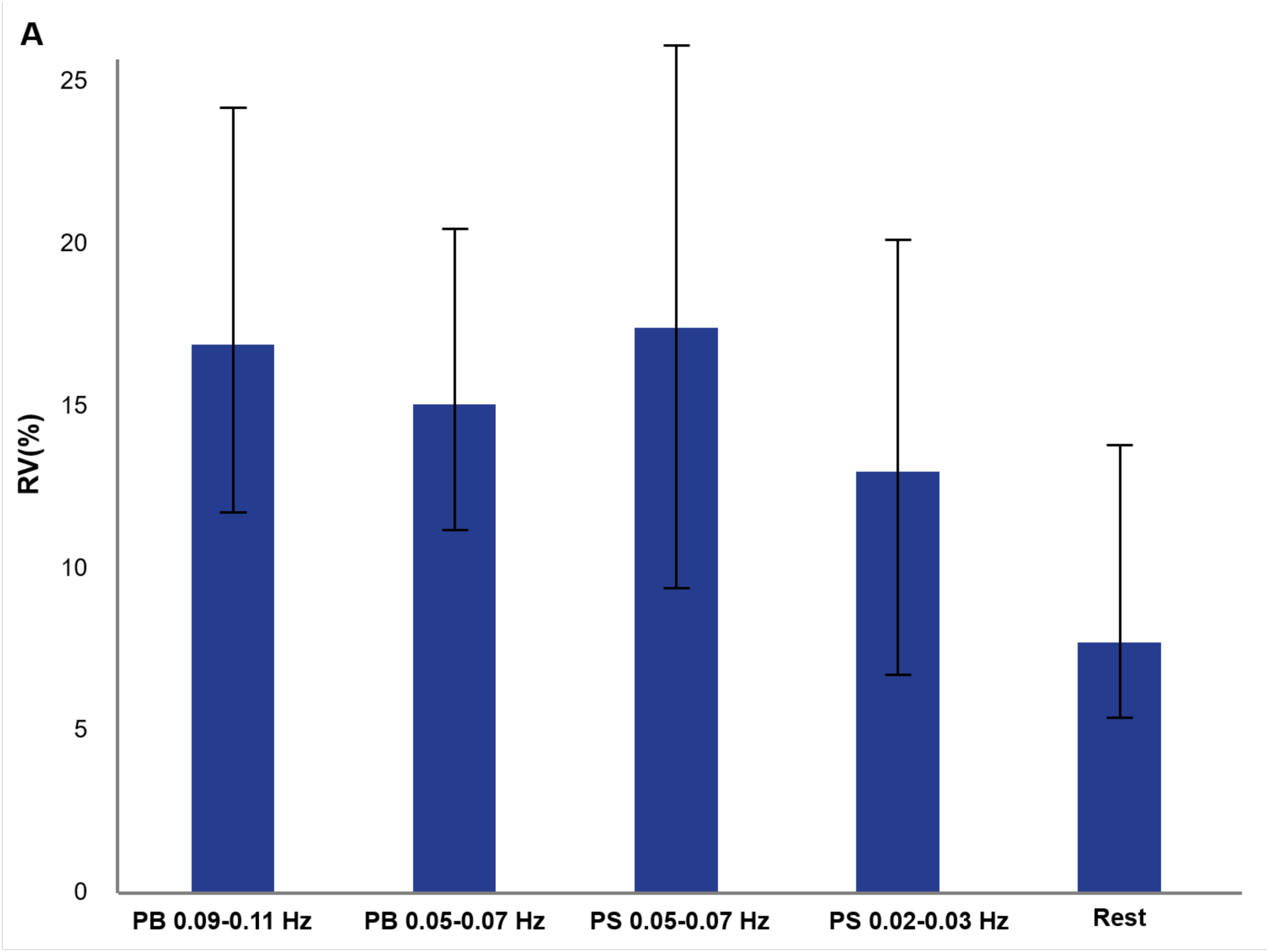
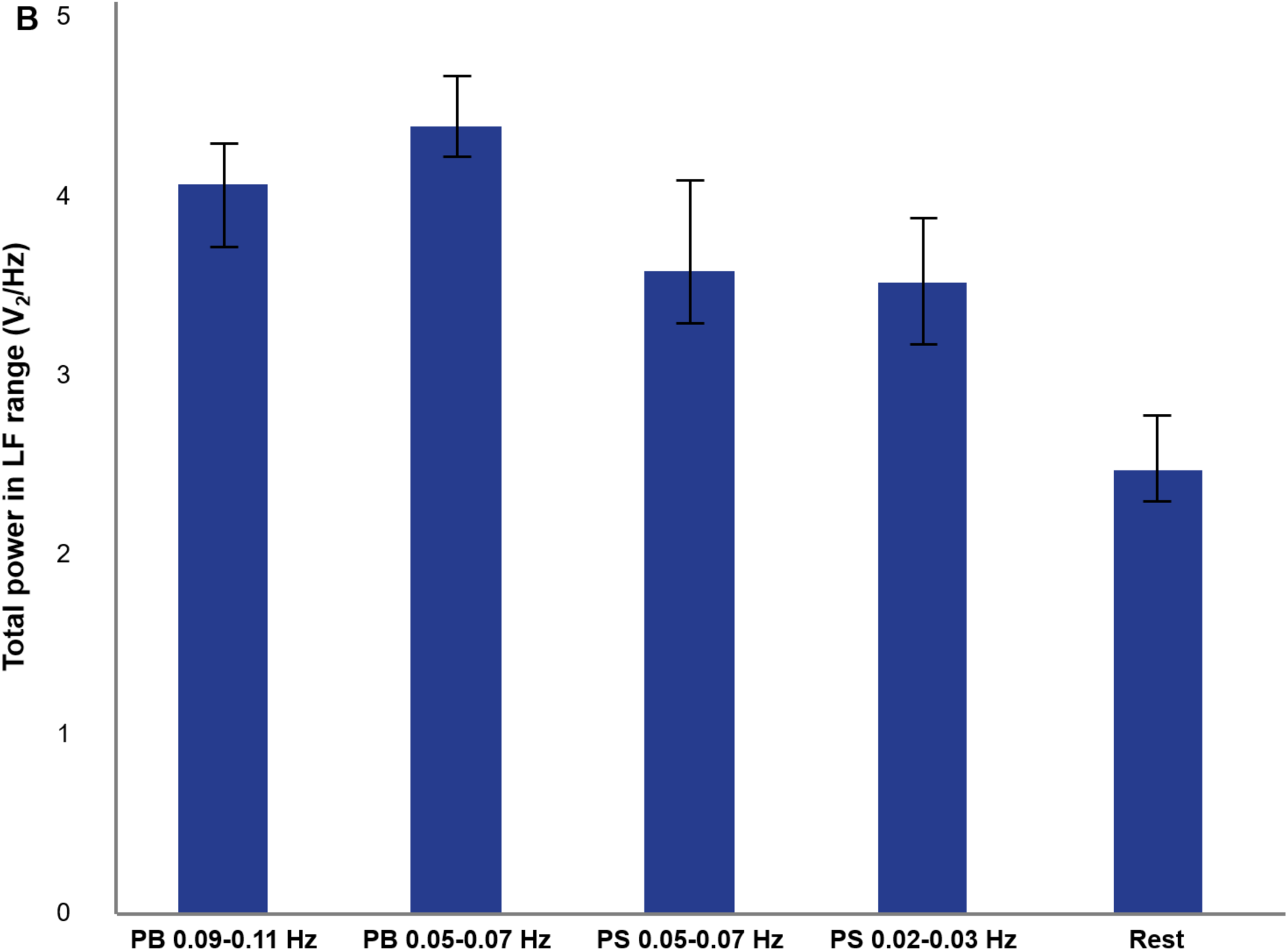
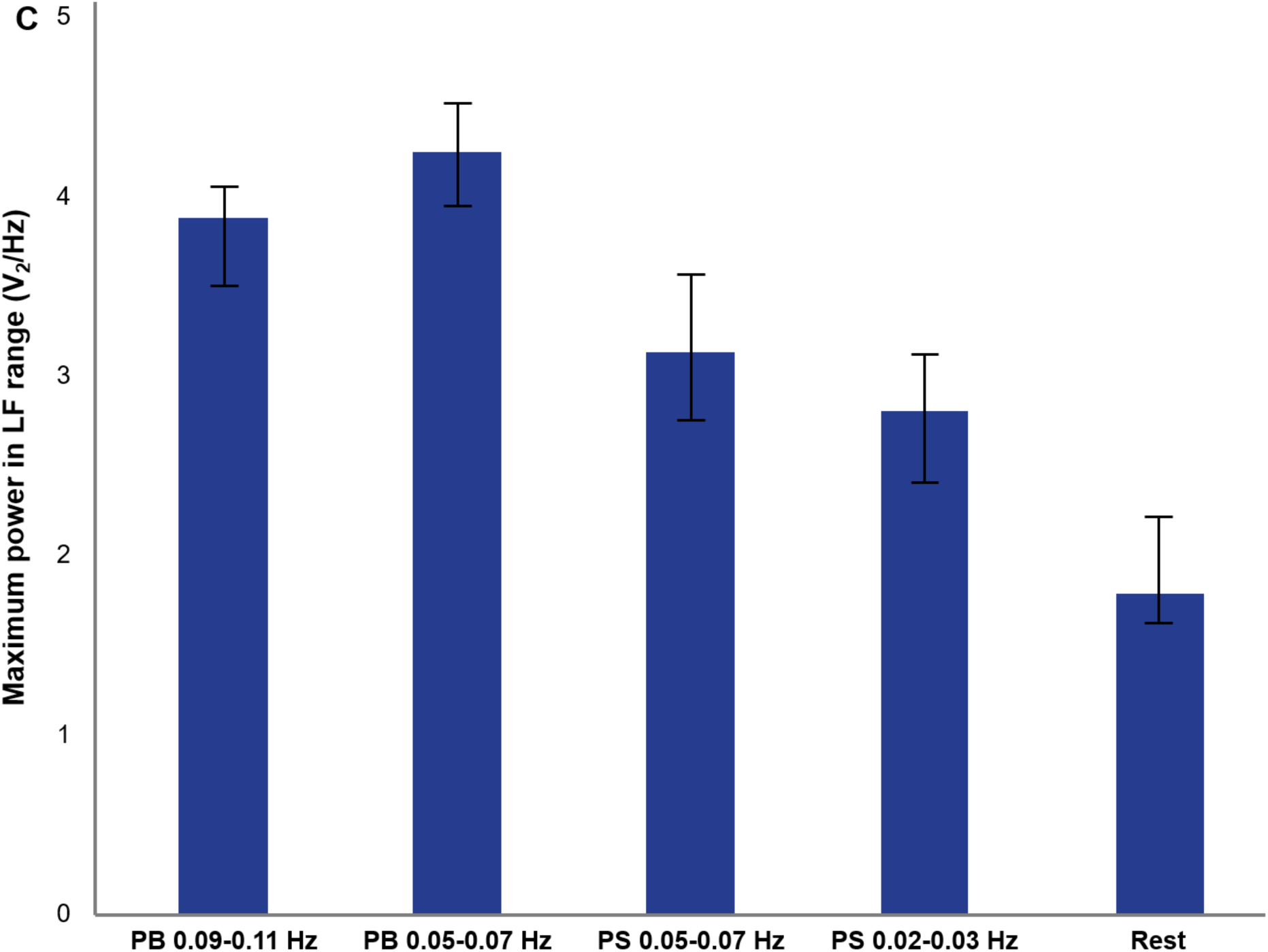
Breathing manipulations significantly change respiration measures compared to rest. (A) median of RV across different conditions. RV is a percentage of inspired volume per second. (B) Median of respiration total power within low frequency range (LF) across different conditions. (C) Median of respiration maximum power within LF range across different conditions. Notes: Error bars are confidence intervals at 95%, PB = paced breathing, PS = paced sighing.

Furthermore, significant pairwise differences between paced sighing at 0.02-0.03 Hz and paced breathing at 0.09-0.011 Hz and paced sighing at 0.05-0.07 Hz were also observed (all FDR-corrected *p*-values < 0.05; Fig. 4A and Supplemental Table 1).

The analysis of the total power of low-frequency oscillations of breathing signals across conditions indicated a significant main effect of condition, *χ*^2^(4) = 105.479, *p*<0.001. Post-hoc pairwise comparisons using the False Discovery Rate (FDR) correction showed that the total power of the breathing signal at rest was significantly lower than the total power of the breathing signal during all the breathing conditions (all FDR-corrected *p*-values < 0.05; Fig. 4B and Supplemental Table 1). Furthermore, analysis of the maximum power of the low-frequency breathing signal across conditions showed a significant main effect of condition, *χ*^2^(4) = 101.455, *p*<0.001. Post-hoc pairwise comparisons using the FDR correction showed that the maximum power of the breathing signal at rest was significantly lower than during all breathing conditions (all FDR-corrected *p*-values < 0.05; Fig. 4C and Supplemental Table 1). Additionally, significant differences were observed among the breathing conditions, with paced breathing at 0.05-0.07 Hz exhibiting the highest maximum and total power (Fig. 3 and Supplemental Table 1).

Furthermore, breathing modulations altered heart rate oscillations. Analysis of PPG data showed that participants’ total heart rate spectral power differed across conditions, *F*(3.31, 105.82) = 13.59, *p* < .001. Post-hoc pairwise comparisons with FDR correction revealed a significant increase in total spectral power during all breathing manipulation scans compared to rest (Fig. 5A). Furthermore, paced sighing at 0.02-0.03 Hz had significantly lower heart rate total spectral frequency power compared to other breathing manipulation conditions (Fig. 5A). In addition, HF, LF, and VLF power each differed significantly across conditions, *F*(3.36, 107.48) = 13.48, *p* < .001; *F*(3.19, 102.10) = 29.65, *p* < .001; and *F*(3.70, 118.53) = 8.11, *p* < .001, respectively. Post-hoc comparisons with FDR correction showed that paced breathing at 0.09–0.11 Hz and 0.05–0.07 Hz significantly decreased HF power compared to rest and paced sighing conditions, while paced sighing at 0.05–0.07 Hz significantly increased HF power relative to rest (all FDR-corrected p < .05; Fig. 5B). Moreover, all breathing manipulations significantly increased LF power compared to rest (all FDR-corrected *p* < .05), and both paced sighing conditions significantly increased VLF power compared to rest (all FDR-corrected *p* < .05; Fig. 5B).

**Fig. 5.**
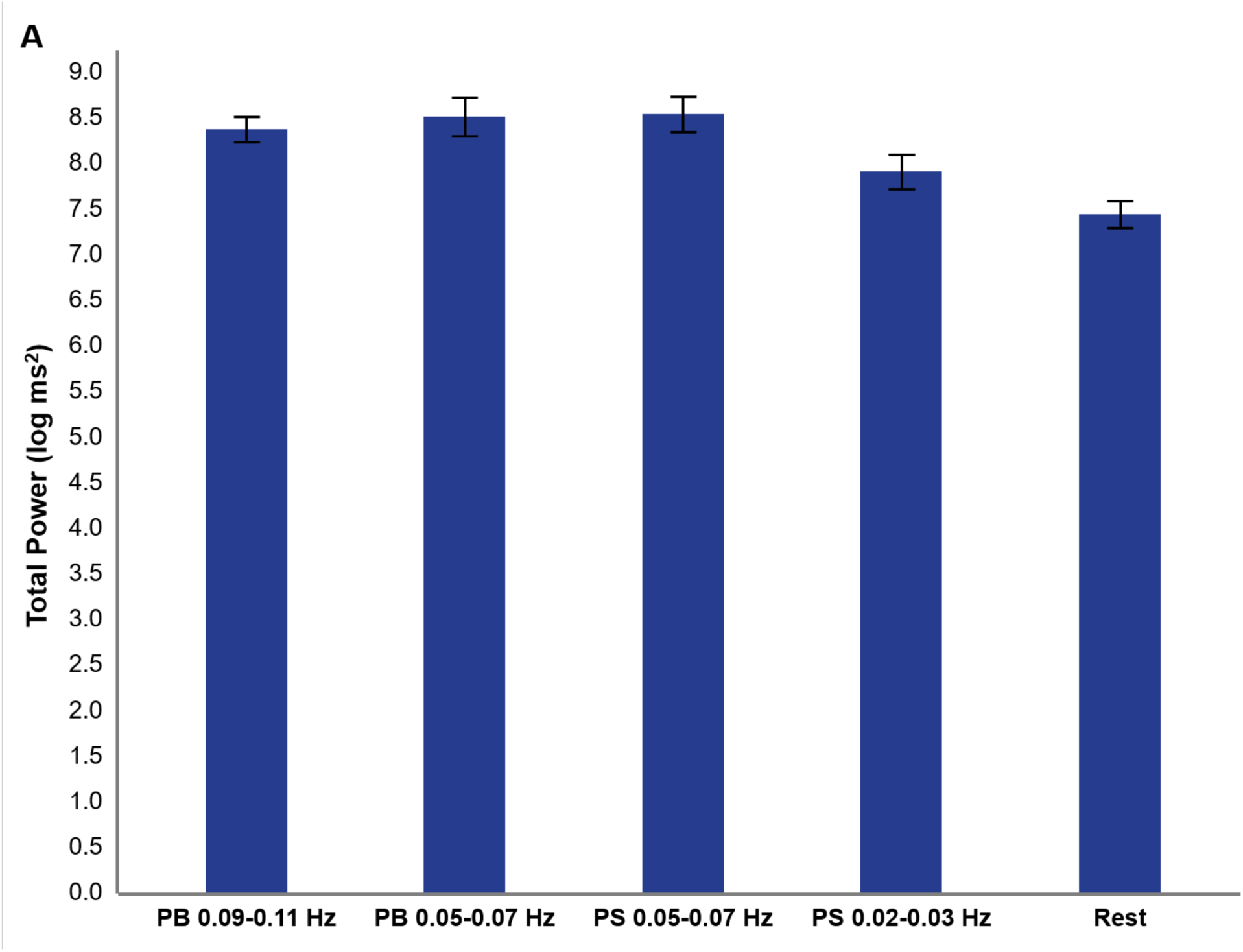
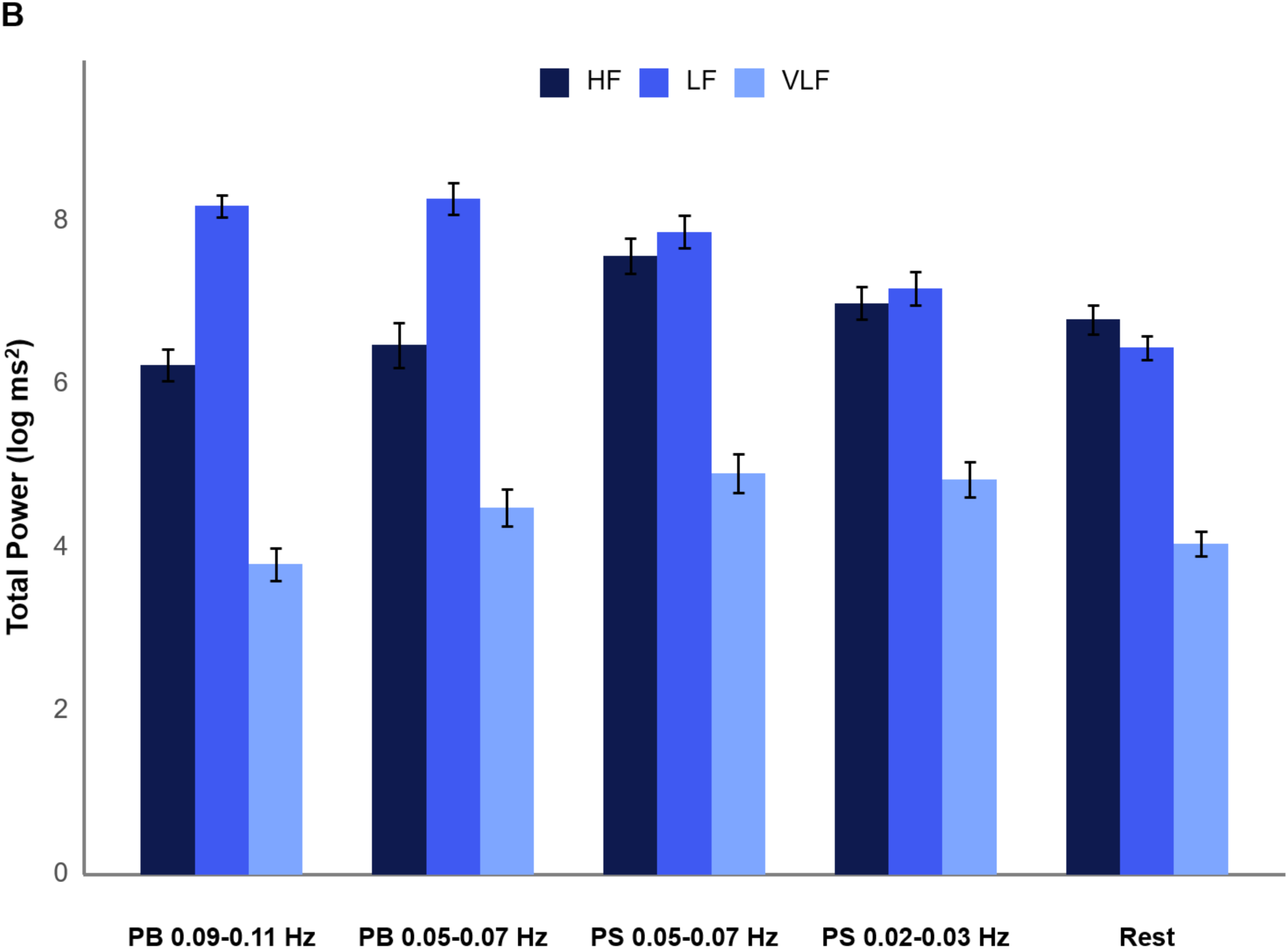
Breathing manipulations significantly change HRV measures compared to rest. (A) mean of log of heart rate total spectral power across different conditions. Error bars are ±1 standard errors. (B) mean of log of high frequency (HF), low frequency (LF), and very low frequency (VLF) powers across different conditions. Notes: Error bars are ±1 standard errors, PB = paced breathing, PS = paced sighing.

### Paced respiration changes CSF spectral power measures

Our breathing manipulation conditions changed the oscillatory patterns of CSF in the fourth ventricle (Fig 6). The repeated measures analysis of low frequency CSF maximum power across conditions revealed a significant main effect of condition, *F*(4,136)=20.063, *p*<0.001. Subsequent post-hoc pairwise comparisons using the FDR correction indicated that the CSF maximum power at rest was significantly lower than in the breathing conditions (Supplemental Table 2 & Fig. 7). No significant differences were observed between the various breathing conditions.

**Fig. 6.**
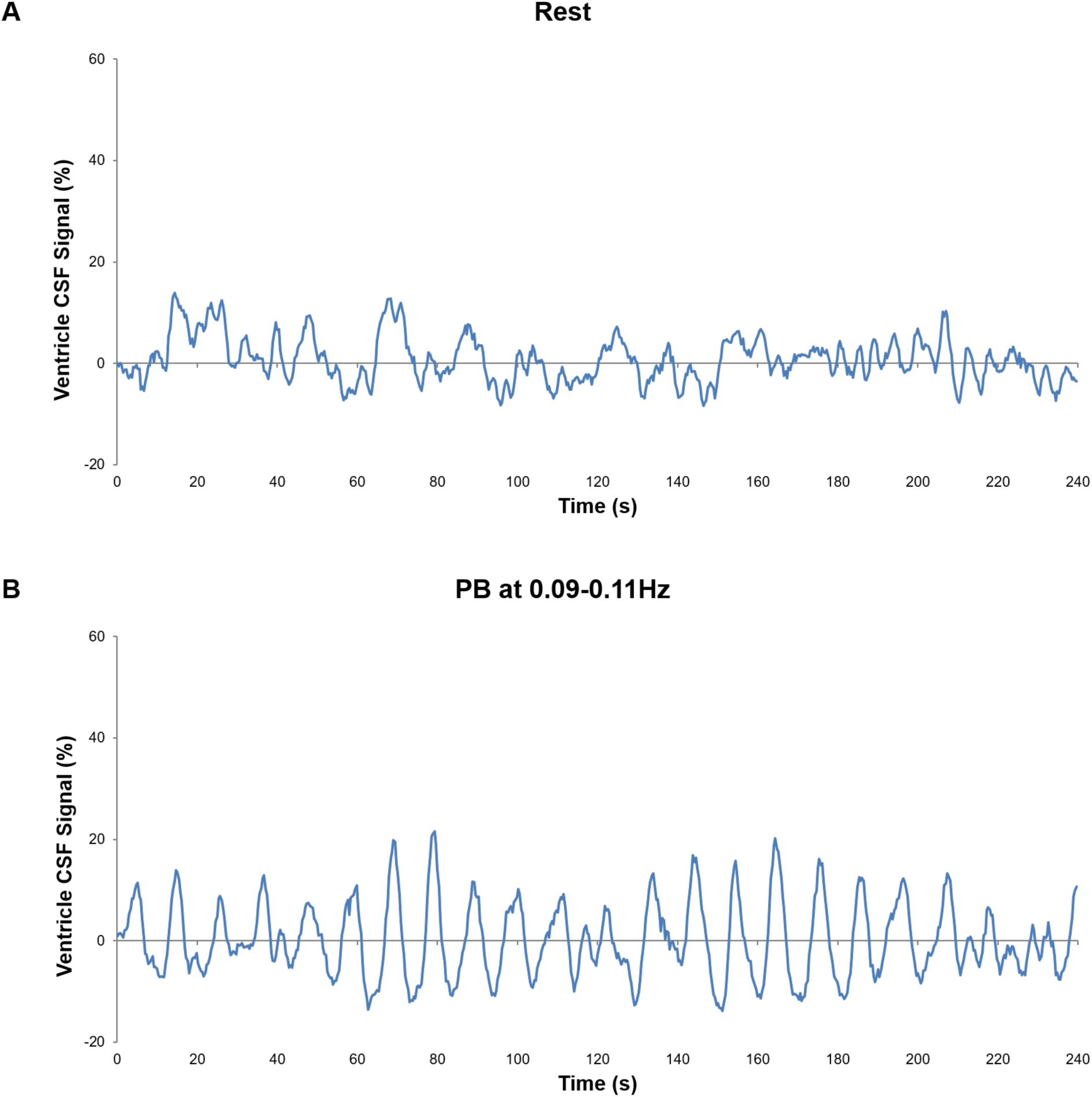
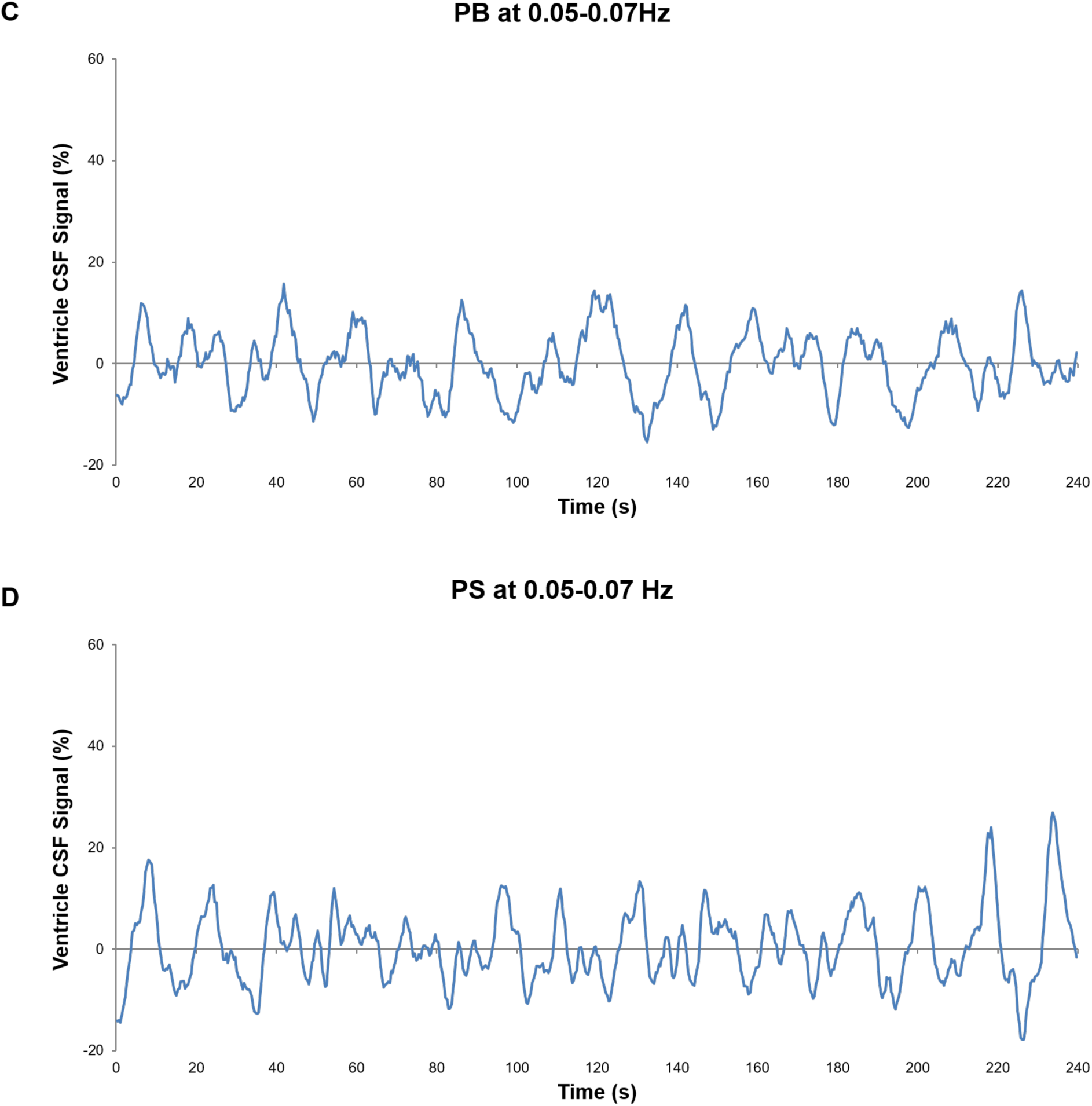
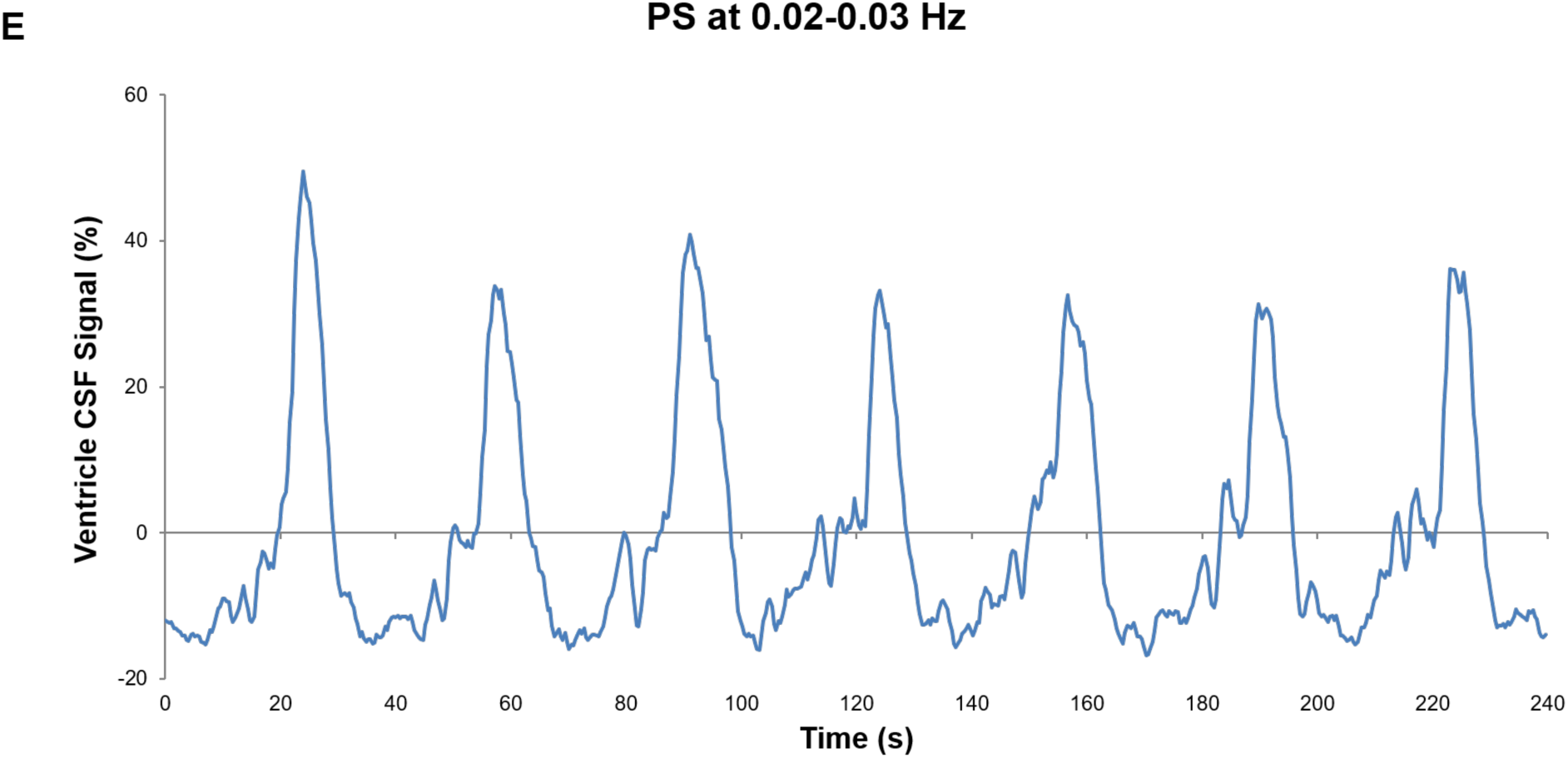
Breathing modulations change CSF oscillations. Example time series of ventricle CSF at (A) rest, (B) paced breathing at 0.09-0.11Hz, (C) paced breathing at 0.05-0.07Hz, (D) paced sighing at 0.05-0.07Hz, and (D) paced sighing at 0.02-0.03Hz in 10 subjects with similar personalized breathing frequencies within frequency ranges of interest. Each participant’s CSF time series is smoothed using a centered 2-second moving average. Note: PB = paced breathing, PS = paced sighing

### Paced sighing at 0.02-0.03 Hz exhibits the highest BOLD global signal amplitude among all conditions

Analysis of BOLD global signal amplitude revealed a significant main effect of condition, *χ*^2^(4) = 28.229, *p*<0.001. Post-hoc pairwise comparisons using the Wilcoxon signed-rank tests with FDR correction showed that all breathing manipulation conditions had significantly higher global signal amplitude than rest (all FDR-corrected *p*-values < 0.05). Additionally, paced sighing at 0.02-0.03 Hz had significantly higher amplitude than paced breathing at 0.09-0.11 Hz (*Z*=2.889, *p*=0.0125) and paced sighing at 0.05-0.07 Hz (*Z*=2.359, *p*=0.030), and a marginally significant higher amplitude than paced breathing at 0.05-0.07 Hz (*Z*=1.982, *p*=0.067) (Fig. 8).

**Fig. 7.**
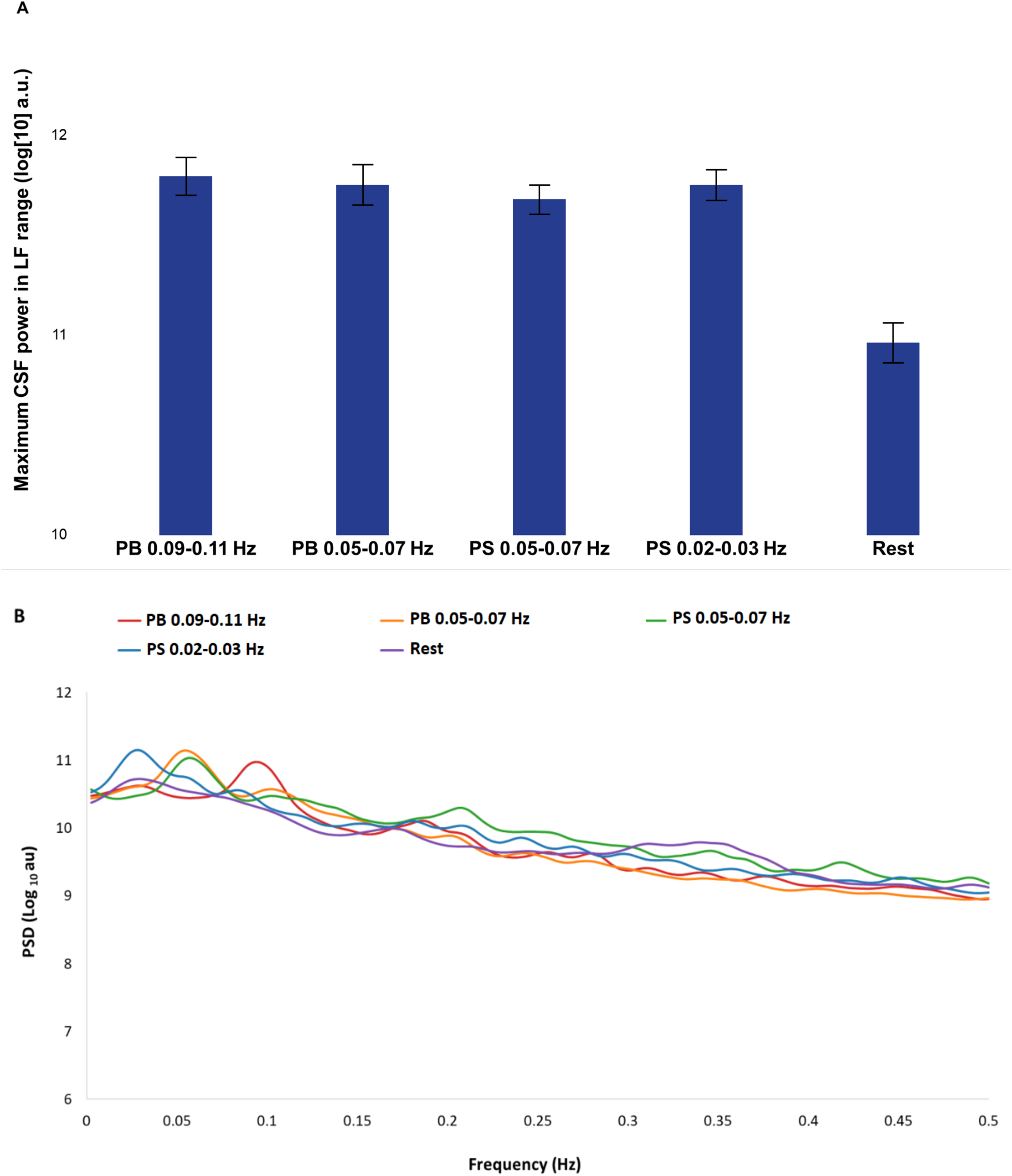
Breathing modulations significantly increase CSF oscillatory power in the low frequency (LF) range. (A) Mean of maximum CSF LF power across different conditions. Error bars are -/+1 standard errors. (B) Power spectral density (PSD) of CSF across different breathing conditions. The data for this graph was further smoothed using a Gaussian-weighted moving average filter. Note: PB = paced breathing, PS = paced sighing.

**Fig. 8.**
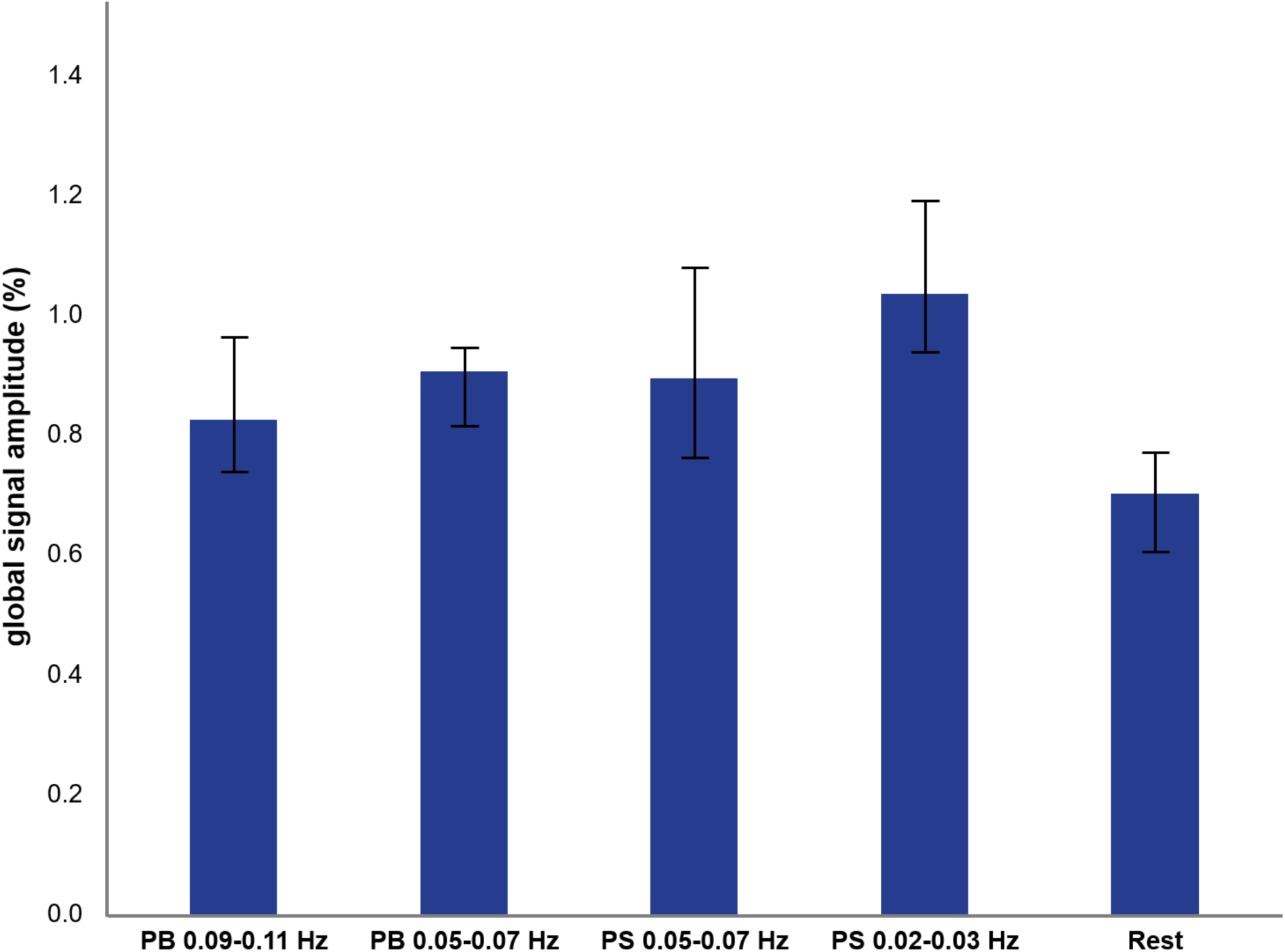
Breathing manipulations significantly increase BOLD signal amplitude compared to rest. Values indicate median of BOLD global signal amplitude. Notes: Error bars are confidence intervals at 95%, PB = paced breathing, PS = paced sighing.

### At rest, zero-thresholded negative derivative of BOLD and CSF shows the strongest correlation at lag 3s

Next, we examined the correlation between CSF and the negative derivative of the BOLD signal during rest. Previous studies have shown that CSF and the negative derivative of the BOLD signal show stronger correlation during NREM sleep compared to wakefulness (Fultz et al., 2019). Analysis of resting state scans revealed a significant correlation between the negative derivative of the BOLD signal and CSF signal. Notably, the peak correlation was observed with a 3-second lag (Monte Carlo *p* < 0.01, cluster-based permutation test, *r* = 0. 0.406), illustrated in Fig. 9. We then investigated correlation patterns under various breathing conditions. Fig. 9 illustrates how these patterns shifted. Across all breathing conditions, significant correlations were found between CSF and the negative derivative of gray matter BOLD signal, albeit with variations in lag and strength. Specifically, paced breathing at 0.09-0.11 Hz, paced breathing at 0.05-0.07 Hz, paced sighing at 0.05-0.07 Hz, and paced sighing at 0.02-0.03 Hz exhibited the strongest correlations at lags -1 s (*r* = 0.285, *p* < 0.01), 1 s (*r* = 0.247, *p* < 0.01), 4 s (*r* = 0.385, *p* < 0.01), and 3 s (*r* = 0.548, *p* < 0.01), respectively.

**Fig. 9.**
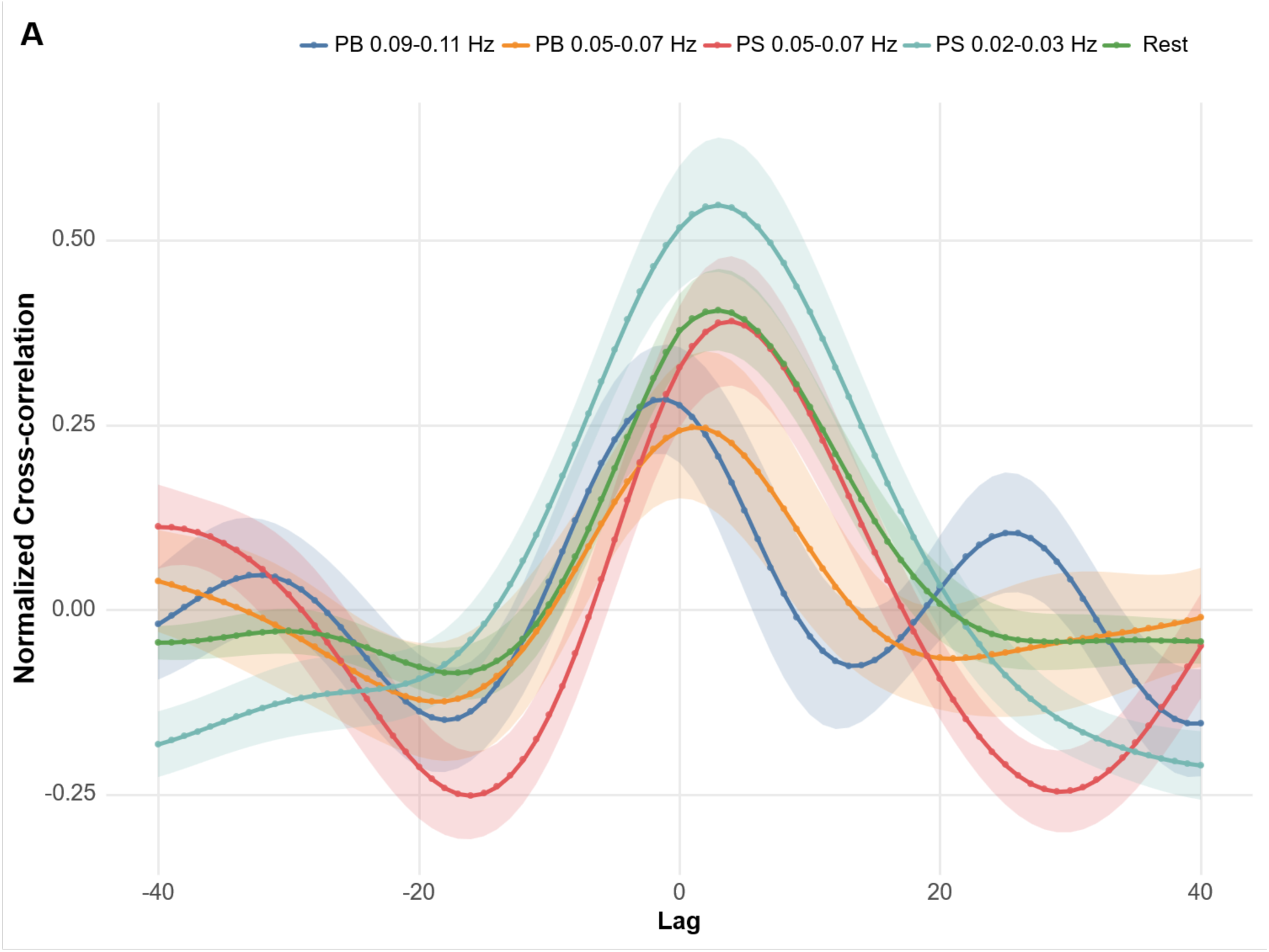
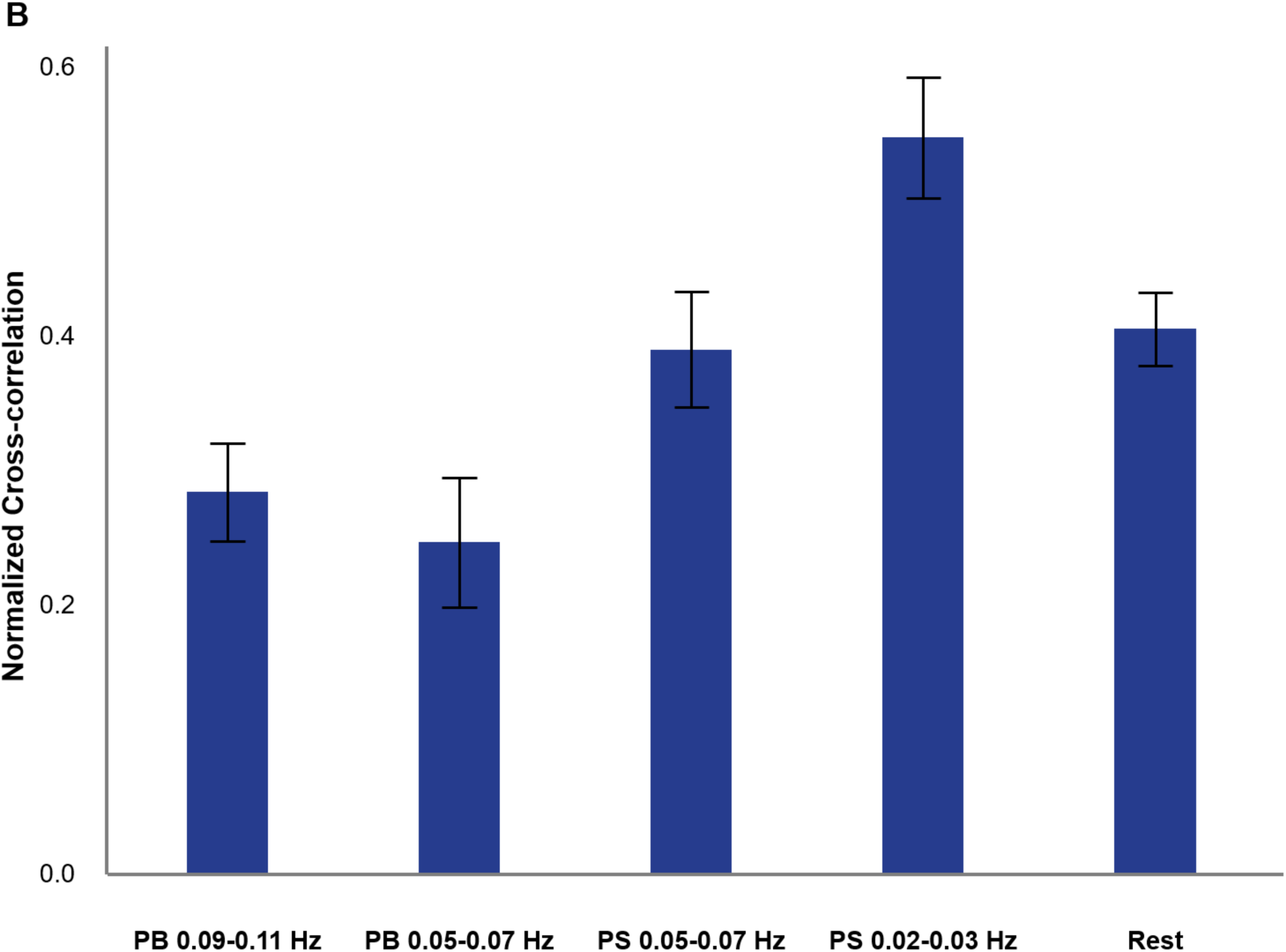
Paced sighing at 0.02-0.03 Hz has the highest cross-correlation of CSF and BOLD signals. (A) The mean cross-correlation between the zero-thresholded negative derivative of BOLD and CSF signals in different conditions. Notes: Shaded areas are 95% confidence intervals. (B) Mean of maximum cross-correlation of CSF and negative derivative of BOLD signal coefficients across different conditions. Notes: Error bars indicate -/+1 standard errors. PB = paced breathing; PS = paced sighing.

We then used repeated measures ANOVA to investigate the effect of breathing conditions on CSF and BOLD signal dynamics. The results revealed a significant main effect of condition on correlation magnitude, *F*(4,136)=13.58, *p*<0.001. Following this significant main effect, post-hoc pairwise comparisons were conducted using Bonferroni correction to identify specific differences between the five conditions. The post-hoc analyses revealed that the CSF and zero-thresholded negative derivatives of the BOLD signal showed the strongest correlation during paced sighing around 0.02-0.03 Hz, which was significantly different from all other conditions (all Bonferroni-corrected *p*-values < 0.05; Fig. 9B). Furthermore, paced breathing at 0.05-0.07 Hz was significantly lower than rest and paced sighing at 0.05-0.07 Hz (all Bonferroni-corrected *p*-values < 0.05).

## Discussion

In the present study, we investigated the effects of four different paced respiratory conditions on measures associated with CSF oscillations. Every condition altered CSF oscillatory power, respiration, cardiac oscillations, and BOLD global-signal amplitude relative to rest. However, only paced sighing at 0.02–0.03 Hz strengthened the coupling between the CSF signal and the negative derivative of the gray-matter BOLD signal, and it did so relative to every other condition, not merely relative to rest.

Firstly, we found that all four paced respiratory conditions increased peak CSF oscillatory power within the low-frequency range (<0.1Hz). This finding is consistent with previous evidence that respiration contributes substantially to CSF dynamics during wakefulness and that controlled breathing can increase CSF pulsatility while shifting CSF and venous flow toward the imposed respiratory frequency (Fultz et al., 2019; Kollmeier et al., 2022).

Furthermore, increased low-frequency CSF power has previously been observed during states of reduced vigilance. During NREM sleep, CSF signals exhibit substantially greater low-frequency oscillatory power than during wakefulness (Fultz et al., 2019). Fourth-ventricle CSF power also increases with drowsiness (Gonzalez-Castillo et al., 2022), and sleep-deprived but awake participants exhibit CSF power comparable to that observed during N1 and N2 sleep (Z. Yang et al., 2025).

Our paced respiratory conditions also increased BOLD global-signal amplitude, with the greatest amplitude observed during paced sighing at 0.02–0.03 Hz. Previous work has consistently associated larger global-signal fluctuations with reduced vigilance. Early investigations reported an increase in global signal amplitude in the visual cortex during early sleep stages (Fukunaga et al., 2006) and a negative correlation with an inverse index of wakefulness in multiple brain regions, including the visual cortex, auditory cortex, and precuneus (Horovitz et al., 2008). Subsequent research established a connection between global signal amplitude and sleep depth (McAvoy et al., 2008), concluding that the rise in global signal amplitude indicates a disproportionately larger decrease in oxygen consumption during sleep relative to the sleep-induced reduction in blood flow. Global signal amplitude is also negatively associated with EEG measures of vigilance during wakefulness, and decreases when vigilance is increased through caffeine administration (Wong et al., 2013). Sleep deprivation similarly increases the amplitude of the global signal (Yeo et al., 2015). The concurrent increase in CSF power and global-signal amplitude therefore suggests that paced respiration, especially paced sighing in the 0.02-0.03 range, may have shifted participants toward a physiological state resembling drowsiness, in which low-frequency cerebrovascular and CSF oscillations become more prominent. This interpretation is consistent with evidence that slow breathing alters neural measures associated with arousal and relaxation (Luo et al., 2025; Sinha et al., 2020; Zaccaro et al., 2018). However, respiration can also affect the global BOLD signal directly due to the induction of vascular dilation and constriction as CO2 levels change; future work tracking multimodal arousal signals could disambiguate these two pathways.

We also examined how the respiratory conditions affected the cross-correlation between the CSF signal and the negative derivative of the BOLD signal. Consistent with the Monro– Kellie relationship and previous imaging studies, CSF fluctuations were strongly coupled to the negative BOLD derivative at rest, reflecting CSF inflow associated with decreases in cerebral blood volume (Fultz et al., 2019; Picchioni et al., 2022; Väyrynen et al., 2026; Yang et al., 2022; Zimmermann et al., 2025). This coupling remained evident under all paced respiratory conditions; but its magnitude varied considerably, and not all changes were increases. Only paced sighing at 0.02–0.03 Hz significantly strengthened the relationship, and it did so relative to every other condition as well as to rest. Paced breathing at 0.05-0.07 Hz, by contrast, produced the weakest coupling observed and fell significantly below rest, with paced breathing at 0.09-0.11 Hz numerically lower than rest as well. Thus, imposing a respiratory oscillation does not necessarily enhance CSF-BOLD coupling and, depending on the frequency, can even weaken it. Previous studies have reported stronger CSF–BOLD coupling during NREM sleep (Fultz et al., 2019; Picchioni et al., 2022), whereas reduced coupling has been observed in age-related diseases such as Alzheimer’s (Han, Chen, et al., 2021; Han et al., 2023), Parkinson’s (Han, Brown, et al., 2021; Wang et al., 2023), and small vessel disease (Zhang et al., 2022). Altered CSF–BOLD coupling has also been associated with greater neocortical tau pathology (Han et al., 2023), higher cortical amyloid-β burden, and subsequent cognitive decline (Han, Chen, et al., 2021).

Notably, the ordering of these coupling effects did not follow the magnitude of the respiratory or cardiac oscillations that the manipulations produced. Paced breathing at 0.05–0.07 Hz generated the highest maximum and total low-frequency respiratory power of any condition, yet yielded the weakest coupling. Nor does it follow cardiac oscillatory power. We selected each participant’s paced frequencies to coincide with peaks in their resting cardiac spectrum, on the expectation that pacing at an intrinsic cardiovascular resonance would amplify cardiac oscillations and, through the Monro–Kellie relationship, CSF oscillations in turn. All manipulations did increase total cardiac spectral power relative to rest, but paced sighing at 0.02–0.03 Hz produced significantly lower total cardiac power than any other manipulation while producing the strongest CSF-BOLD coupling. Taken together, these dissociations indicate that the effect depends on the frequency of the imposed oscillation rather than on the amplitude of the respiratory or cardiovascular response it evokes.

The enhanced CSF–BOLD coupling observed during paced sighing at 0.02–0.03 Hz is consistent with growing evidence that infraslow vascular and autonomic oscillations contribute to CSF movement. Paced sighing at this frequency produced the largest increase in global-signal amplitude and significantly strengthened CSF–BOLD coupling relative to rest, suggesting that respiratory oscillations imposed within this band may engage vascular dynamics similar to those observed during sleep in rodents (Hauglund et al., 2025) and humans (Väyrynen et al., 2026).

One possible mechanism is that repeated sighing at 0.02–0.03 Hz amplified infraslow fluctuations in vascular tone and blood pressure. A sigh consists of a rapid, deep inspiration followed by a slower expiration and can produce a transient reduction in vascular tone and blood pressure, followed by compensatory changes in heart rate and a gradual return toward baseline (E. G. Vaschillo et al., 2015; Wilhelm et al., 2001). When sighs are repeated at regular intervals, these responses can produce pronounced low-frequency cardiovascular oscillations, and may therefore influence afferent outflow from baroreceptors and enhance neural inhibition (B. Vaschillo & Vaschillo, 2020; E. G. Vaschillo et al., 2015). The photoplethysmographic intensity ratio, an index sensitive to slow blood-pressure and peripheral vascular-tone changes, shows particularly strong coherence with CSF signals at approximately 0.02–0.03 Hz (Attarpour et al., 2021; Ding et al., 2017). Repeated sighing within this band may therefore have amplified vascular-volume changes at a frequency to which the CSF signal is especially responsive. This interpretation is also consistent with the difference between the two sighing conditions. Both involved repeated sighs, but only sighing at 0.02–0.03 Hz significantly increased CSF–BOLD coupling. The effect is therefore unlikely to reflect sighing alone. Instead, it may depend on the interval at which sigh-related cardiovascular responses recur. However, because the 0.02–0.03 Hz condition was implemented only through paced sighing, the effects of respiratory frequency and respiratory form cannot be fully separated. A manipulation capable of generating sustained respiratory or vascular oscillations at 0.02–0.03 Hz without repeated sighs would be needed to determine whether the enhanced coupling resulted from the infraslow frequency itself, the physiological characteristics of sighing, or their combination.

Changes in arterial CO₂ provide another plausible mechanism. Variations in arterial CO₂ pressure alter vascular tone via the arterial baroreflex (Ainslie et al., 2005; Badra et al., 2001; Harris et al., 2006), with hypercapnia producing vasodilation and increased cerebral blood flow and hypocapnia producing vasoconstriction and reduced cerebral blood flow (Aaslid et al., 1989; Nair et al., 2024). End-tidal CO₂ fluctuations during rate-controlled resting-state breathing are associated with global BOLD fluctuations, including prominent effects at approximately 0.03 Hz (Birn et al., 2006). Deep sighs and prolonged exhalations may produce larger or more slowly varying changes in CO₂ than ordinary paced breathing. These fluctuations could increase the amplitude and temporal regularity of cerebral blood-volume changes and thereby strengthen their coupling with CSF displacement.

The longer expiration characteristic of the sighing conditions may also have contributed. Extending exhalation relative to inhalation is thought to promote greater physiological and psychological relaxation, and increasing the exhalation-to-inhalation ratio leads to stronger activation of parasympathetic activity (Birdee et al., 2023). Our own data suggest, however, that extended exhalation cannot be sufficient on its own. Both sighing conditions used the same 1 s inhalation followed by 3 s exhalation, and the only condition to significantly increase high-frequency HRV relative to rest was the sighing condition that did not enhance coupling (paced sighing at 0.05–0.07 Hz), whereas both paced breathing conditions significantly reduced HF power. Parasympathetic engagement and coupling strength therefore dissociate in this dataset. Future studies are needed to assess how paced sighing affects cardiovascular and CSF-related measures compared to paced breathing with extended exhalation.

Several limitations of our study should be considered. First, we did not measure end-tidal CO₂ or continuous blood pressure, preventing us from determining how changes in vascular tone, arterial pressure, and respiratory gas exchange contributed to the BOLD and CSF signals oscillations. Second, we did not collect direct measures of central arousal or noradrenergic activity, such as simultaneous EEG, or pupillometry. The interpretation that paced sighing engaged low-arousal or noradrenergically mediated dynamics therefore remains provisional. Third, the present CSF measurements were obtained from the fourth ventricle and therefore primarily reflect bulk CSF displacement within the ventricular system. Our findings therefore identify a plausible hemodynamic driver of CSF movement but provide only indirect evidence regarding fluid dynamics within the interstitial and perivascular compartments. CSF-STREAM would provide complementary information at a smaller spatial scale by measuring CSF mobility across ventricular, subarachnoid, and perivascular compartments (Hirschler et al., 2025). This approach could test whether the increased ventricular CSF–BOLD coupling observed during paced sighing is accompanied by greater CSF mobility around penetrating vessels, closer to the sites at which CSF may interact with interstitial fluid (Hirschler et al., 2025). Additionally interleaving fast fMRI acquisition with real-time phase contrast MRI measurement in the fourth ventricle can elucidate more information on effect of paced respiratory oscillations on CSF movement by allowing to quantify real-time CSF flow dynamics along with global BOLD fluctuations (Roefs et al., 2024).

Respiration is vital for maintaining homeostasis and adapting to environmental demands. Furthermore, it can act as a gateway to our physiology. By modulating our breathing frequency, we can influence physiological mechanisms associated with it, including cardiac pulsations and the brain’s mechanisms such as CSF movement. Our study explored how different breathing frequencies and paced breathing versus paced sighing affect CSF movement into the brain and the mechanisms associated with increased CSF flow, and found that which frequency is chosen matters more than how vigorously the manipulation is performed: paced sighing at 0.02–0.03 Hz strengthened the coupling between cerebral blood volume and CSF displacement beyond every other condition tested, while the condition generating the strongest respiratory oscillation weakened it. Further research is necessary to examine how various physiological parameters, including respiration, heart rate, blood pressure, and peripheral vascular tone, influence CSF movement under different breathing conditions, how these effects are modulated by aging, and whether engaging this frequency band during wakefulness translates into enhanced clearance.

## Acknowledgements

This research was supported by National Institutes of Health grants R01AG025340, R01AG057184, R01AG080652, and R01AT011429.

## Statements and Declarations

### Funding

This research was supported by R01AG025340, R01AG057184, R01AG080652, and R01AT011429.

### Competing Interests

The authors have no relevant financial or non-financial interests to disclose.

### Author Contributions

P.N.: Conceptualization, Methodology, Investigation, Data Curation, Formal analysis, Project Administration, Software, Writing—original draft, Writing—review and editing, and Visualization. M.B.: Data Curation, Formal Analysis, Validation, Software, and Writing—review and editing. A.L.: Investigation, Data Curation, Writing—review and editing. H.Y.: Methodology, Software, Writing—review and editing. L.L: Methodology, Conceptualization, Writing—review and editing. M.M.: Funding Acquisition, Conceptualization, Methodology, Supervision, Resources, and Writing—review and editing.

### Data Availability

Data from this project will be made available on OpenNeuro.org upon publication.

### Ethics Approval

All procedures performed in this study were approved by the USC Institutional Review Board (UP-20-01022).

## Supplementary Materials

**Supplemental Table 1.**
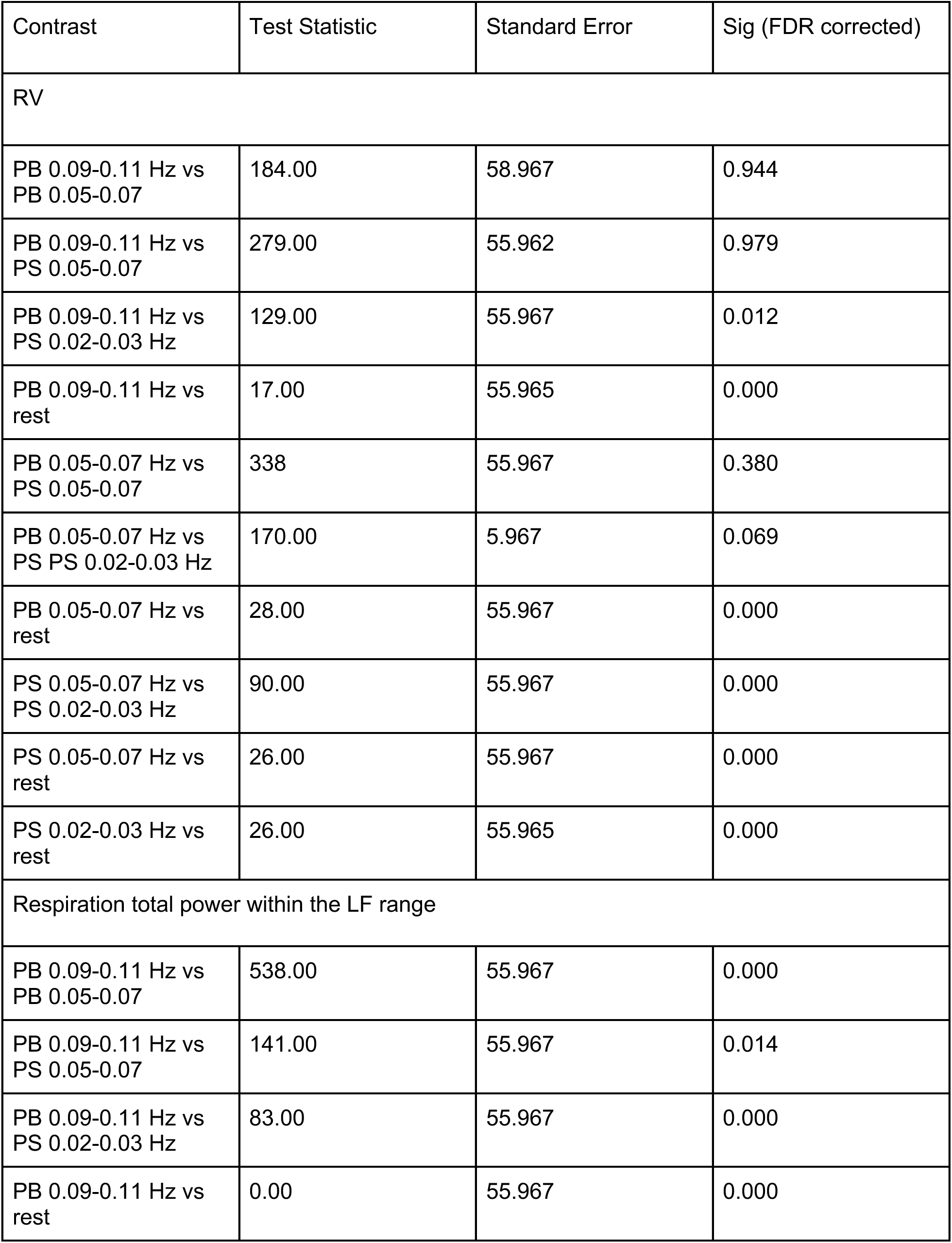
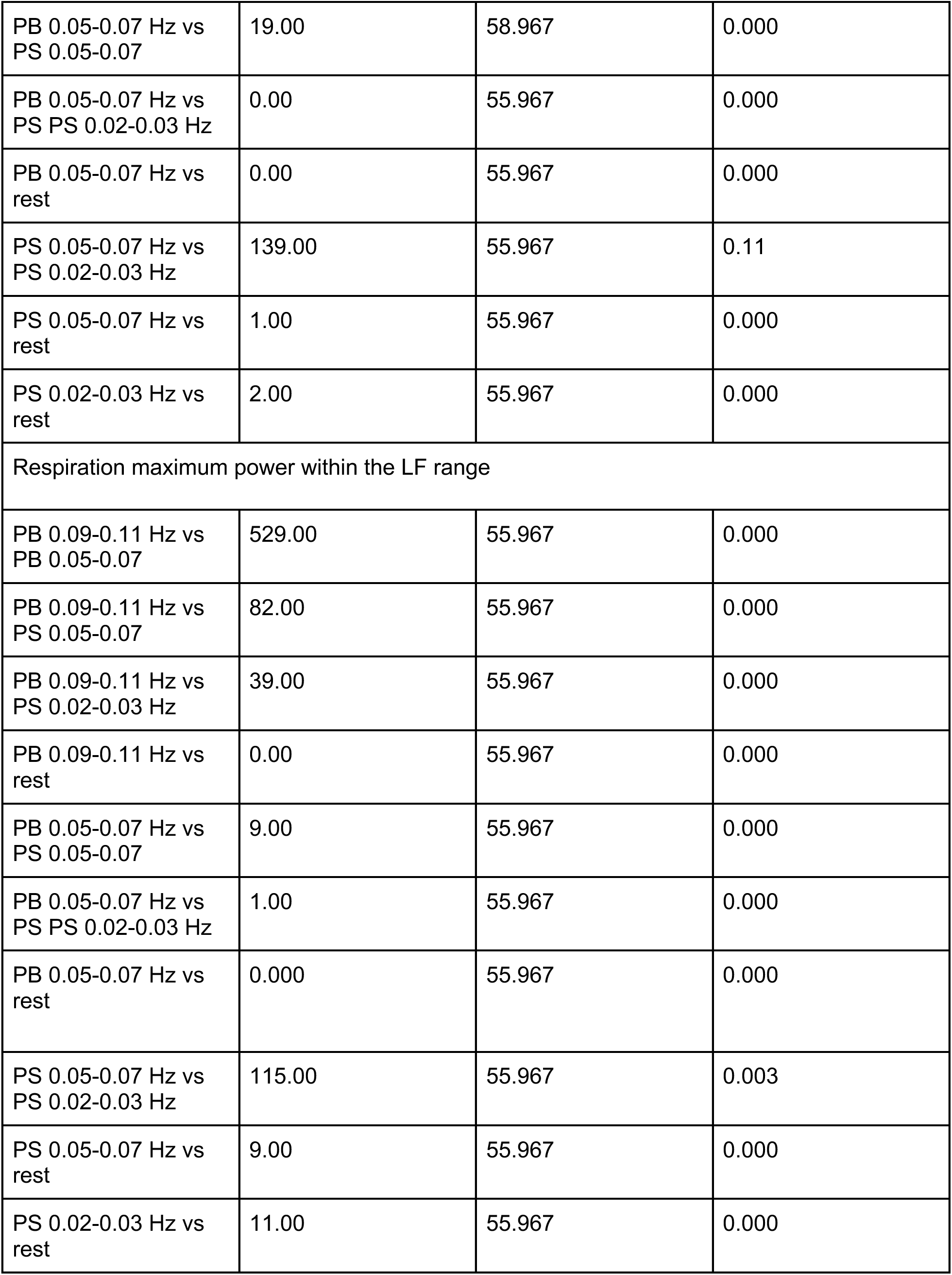
Results of post-hoc FDR-corrected pairwise comparisons of respiratory measures (RV, Total power within the low frequency range, and maximum power within the low frequency range). All breathing manipulation conditions had significantly higher RV, total power, and maximum power compared to rest.

**Supplemental Table 2.**
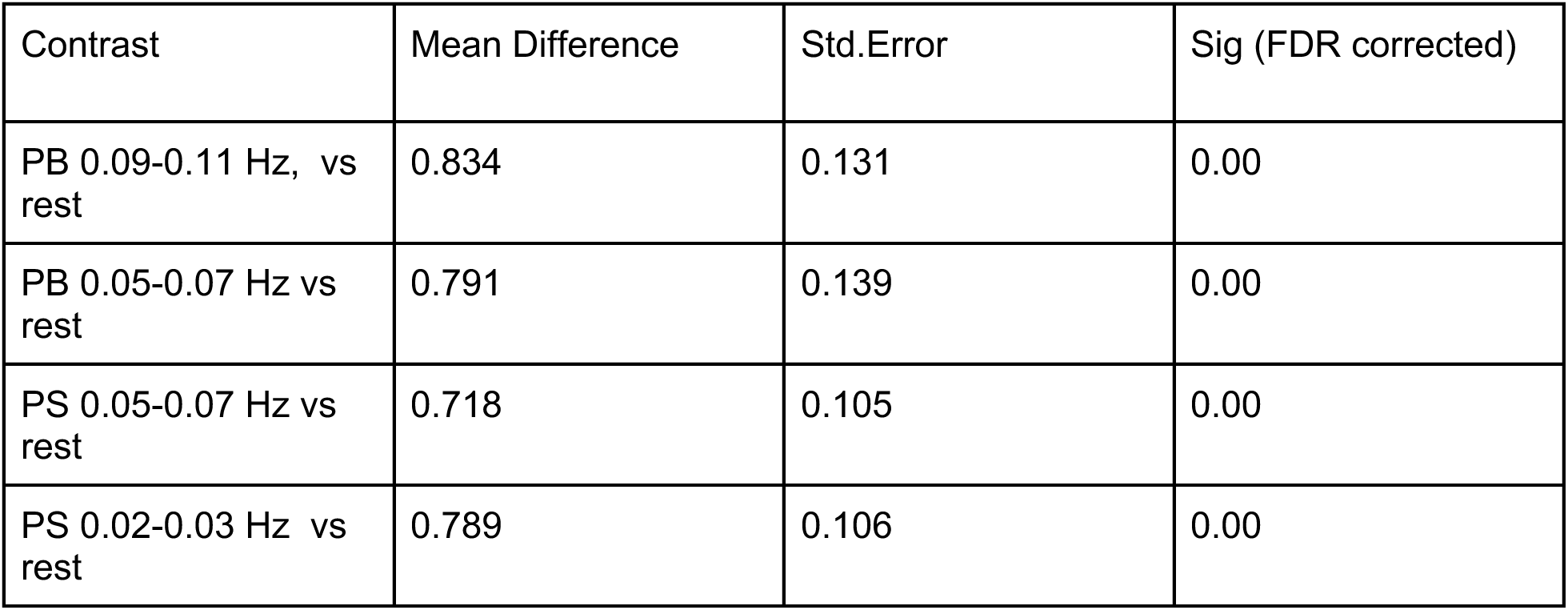
Results of post-hoc FDR-corrected pairwise comparisons of CSF maximum power during different conditions. Breathing manipulations significantly change CSF maximum power compared to rest. No significant differences were found between breathing manipulation conditions.

## Notes

### Competing Interest Statement

The authors have declared no competing interest.

### Summary of Updates

The manuscript has been substantially revised including a rewritten Introduction and revisions to the Discussion, to more clearly articulate the rationale for the study, the selection of specific breathing frequencies and modulation types, and the interpretation of our findings. We have also expanded the discussion of the limitations. In addition, we now provide a more detailed description of the breathing conditions and the instructions given to participants.

