## Supplementary Materials for "Effects of different slow paced breathing regimes on cerebrospinal fluid (CSF) oscillations"

**Supplemental Table 1**

Results of post-hoc FDR-corrected pairwise comparisons of respiratory measures (RV, Total power within the low frequency range, and maximum power within the low frequency range). All breathing manipulation conditions had significantly higher RV, total power, and maximum power compared to rest.

| Contrast | Test Statistic | Standard Error | Sig (FDR corrected) |
| --- | --- | --- | --- |
| RV | | | |
| PB 0.09-0.11 Hz vs PB 0.05-0.07 | 184.00 | 58.967 | 0.944 |
| PB 0.09-0.11 Hz vs PS 0.05-0.07 | 279.00 | 55.962 | 0.979 |
| PB 0.09-0.11 Hz vs PS 0.02-0.03 Hz | 129.00 | 55.967 | 0.012 |
| PB 0.09-0.11 Hz vs rest | 17.00 | 55.965 | 0.000 |
| PB 0.05-0.07 Hz vs PS 0.05-0.07 | 338 | 55.967 | 0.380 |
| PB 0.05-0.07 Hz vs PS PS 0.02-0.03 Hz | 170.00 | 5.967 | 0.069 |
| PB 0.05-0.07 Hz vs rest | 28.00 | 55.967 | 0.000 |
| PS 0.05-0.07 Hz vs PS 0.02-0.03 Hz | 90.00 | 55.967 | 0.000 |
| PS 0.05-0.07 Hz vs rest | 26.00 | 55.967 | 0.000 |
| PS 0.02-0.03 Hz vs rest | 26.00 | 55.965 | 0.000 |
| Respiration total power within the LF range | | | |
| PB 0.09-0.11 Hz vs PB 0.05-0.07 | 538.00 | 55.967 | 0.000 |
| PB 0.09-0.11 Hz vs PS 0.05-0.07 | 141.00 | 55.967 | 0.014 |
| PB 0.09-0.11 Hz vs PS 0.02-0.03 Hz | 83.00 | 55.967 | 0.000 |
| PB 0.09-0.11 Hz vs rest | 0.00 | 55.967 | 0.000 |
| PB 0.05-0.07 Hz vs PS 0.05-0.07 | 19.00 | 58.967 | 0.000 |
| PB 0.05-0.07 Hz vs PS PS 0.02-0.03 Hz | 0.00 | 55.967 | 0.000 |
| PB 0.05-0.07 Hz vs rest | 0.00 | 55.967 | 0.000 |
| PS 0.05-0.07 Hz vs PS 0.02-0.03 Hz | 139.00 | 55.967 | 0.11 |
| PS 0.05-0.07 Hz vs rest | 1.00 | 55.967 | 0.000 |
| PS 0.02-0.03 Hz vs rest | 2.00 | 55.967 | 0.000 |
| Respiration maximum power within the LF range | | | |
| PB 0.09-0.11 Hz vs PB 0.05-0.07 | 529.00 | 55.967 | 0.000 |
| PB 0.09-0.11 Hz vs PS 0.05-0.07 | 82.00 | 55.967 | 0.000 |
| PB 0.09-0.11 Hz vs PS 0.02-0.03 Hz | 39.00 | 55.967 | 0.000 |
| PB 0.09-0.11 Hz vs rest | 0.00 | 55.967 | 0.000 |
| PB 0.05-0.07 Hz vs PS 0.05-0.07 | 9.00 | 55.967 | 0.000 |
| PB 0.05-0.07 Hz vs PS PS 0.02-0.03 Hz | 1.00 | 55.967 | 0.000 |
| PB 0.05-0.07 Hz vs rest | 0.000 | 55.967 | 0.000 |
| PS 0.05-0.07 Hz vs PS 0.02-0.03 Hz | 115.00 | 55.967 | 0.003 |
| PS 0.05-0.07 Hz vs rest | 9.00 | 55.967 | 0.000 |
| PS 0.02-0.03 Hz vs rest | 11.00 | 55.967 | 0.000 |

**Supplemental Table 2**

Results of post-hoc FDR-corrected pairwise comparisons of CSF maximum power during different conditions. Breathing manipulations significantly change CSF maximum power compared to rest. No significant differences were found between breathing manipulation conditions.

| Contrast | Mean Difference | Std.Error | Sig (FDR corrected) |
| --- | --- | --- | --- |
| PB 0.09-0.11 Hz, vs rest | 0.834 | 0.131 | 0.00 |
| PB 0.05-0.07 Hz vs rest | 0.791 | 0.139 | 0.00 |
| PS 0.05-0.07 Hz vs rest | 0.718 | 0.105 | 0.00 |
| PS 0.02-0.03 Hz vs rest | 0.789 | 0.106 | 0.00 |
